# CircZBTB46, a promising therapeutic target in crizotinib resistant ALK-positive T lymphomas

**DOI:** 10.64898/2026.01.13.699235

**Authors:** Loélia Babin, Elissa Andraos, Steffen Fuchs, Chloe Bessière, Lola Colras, Cathy Quelen, Cindy Pinto, Iyad Daoudi, Romain Pfeifer, Ahmed Zamani, Marina Bousquet, Stéphane Pyronnet, Christine Gaspin, Laurence Lamant, Fabienne Meggetto

**Author notes:** Correspondance: Fabienne MEGGETTO CRCT, 2 Avenue Hubert Curien 31037 Toulouse, France, Phone: (+)33 (5)62-74-45-39, Fax: (+)33 (5)62-74-45-58. For original data, please contact.

## Abstract

Circular RNAs (circRNAs) are increasingly recognized as functional non-coding transcripts with oncogenic potential. Here, a comprehensive analysis of circRNA expression in primary ALK(+) anaplastic large-cell lymphoma (ALK(+) ALCL) is presented. Integrated transcriptomic profiling revealed that aberrant expression of circZBTB46 and of its linear host transcript, normally restricted to dendritic cells, is exclusive to ALK(+) lymphoma cells and driven by the oncogenic NPM1-ALK/STAT3 axis. Functional studies showed that circZBTB46, unlike its protein-coding counterpart, promotes resistance to the ALK inhibitor crizotinib. Silencing circZBTB46 restored crizotinib sensitivity in resistant ALCL cells both *in vitro* and *in vivo*. Transcriptomic analyses identified PIP5K1C as a downstream effector regulated through a competitive endogenous RNA mechanism in which circZBTB46 acts as a sponge to miR-25-3p, alleviating its repression of PIP5K1C. These findings uncover a previously unrecognized mechanism of drug resistance in ALK(+) ALCL and establish circZBTB46 as a promising therapeutic target.

## Introduction

The purpose of this study was to investigate circRNAs in the specific context of ALK(+) ALCL. Anaplastic large-cell lymphoma (ALCL) is a rare and aggressive peripheral T-cell non-Hodgkin lymphoma (NHL) belonging to CD30^+^ lymphoproliferative disorders^1^. ALCL is divided into two subtypes based on the expression of the anaplastic lymphoma kinase (ALK)^2^. The ALK(+) subtype primarily affects children and young adults. In approximately 80–85% of ALK(+) cases, the t(2;5)(p23;q35) chromosomal translocation is present, driving the fusion of the intracytoplasmic domain of ALK, on chromosome 2p23, with the N-terminal portion of nucleophosmin (NPM1), on chromosome 5q35^3^. This NPM1-ALK fusion protein constitutively activates key oncogenic signaling pathways, including RAS-ERK, JAK3-STAT3 and PI3K-Akt, thereby driving cancer cell proliferation, differentiation and survival^4^. In ALK(+) ALCL, current first-line multidrug chemotherapy (CT) regimens achieve a progression-free survival of approximately 70% at ten years after diagnosis. Crizotinib, the first ALK-targeted tyrosine kinase inhibitor (TKI), was introduced for the treatment of ALK(+) ALCL refractory or relapsing (R/R) after CT. However, the clinical efficacy of crizotinib has been hampered by the emergence of resistance. The prognosis for R/R patients remains poor, highlighting the need for a better understanding of the aggressive behavior of ALCL and the development of novel prognostic markers to identify high-risk patients at an early stage^5,6^. Previous studies have disclosed the important role of small noncoding RNAs, particularly microRNAs (miRNAs), in ALCL^7^. We and others have shown that specific miRNAs are associated with an increased risk of relapse in ALK(+) ALCL, after CT or crizotinib treatment^8–13^. Circular RNAs (circRNAs) are a recently recognized class of stable and conserved noncoding RNAs characterized by a covalently closed loop structure lacking 5′ and 3′ ends. Predominantly localized in the cytoplasm, circRNAs act as molecular sponges for microRNAs and RNA-binding proteins, thereby modulating post-transcriptional gene regulation. They are highly stable and often exhibit tissue- or stage-specific expression patterns. Dysregulation of circRNAs has been shown to be involved in the pathophysiology of several solid tumors and hematological malignancies, especially B-NHL and leukemias^14,15^. A few reports have addressed the implication of circRNAs in peripheral T-cell NHL such as T-cell lymphoblastic lymphoma^15,16^, but to date, no studies appear to have examined their role in ALK(+) ALCL. Deciphering the functions and mechanisms of circRNAs in ALK(+) ALCL could lead to the identification of innovative biomarkers development of therapeutic

## Results

### Aberrant expression of circRNA and mRNA *ZBTB46* in NPM1-ALK(+) lymphoma cells

This work was initiated by investigating genome-wide circRNA expression profiles in ALK(+) ALCL via various approaches.

First, a ribo-minus RNA sequencing (RNAseq) dataset, previously generated from a cohort of 39 ALK(+) ALCL biopsies, was analyzed, with 9 reactive lymph nodes (RLNs) as controls^17^. A circRNA expression landscape was established in ALK(+) ALCL (log2Fold Change ≥ 2 and a *P* value <0.05, **Fig. 1A**). Among upregulated circRNAs, circZBTB46 displayed the highest expression (L2FC: 3.6; **Fig. 1A and 1B**). This circRNA stood out for several reasons. First, using long-read Oxford Nanopore RNA-seq, our group has already identified circZBTB46 as one of the most abundant circRNAs expressed in four human ALK(+) ALCL cell lines^18^. Second, the expression of its host gene, *ZBTB46*, is known to be restricted to human progenitor and conventional dendritic cells but has not been reported in T-lymphocytes^19,20^. Finally, using the RNAseq dataset, it was found that both *ZBTB46* circRNA and mRNA were undetectable in RLN (**Fig. 1C**).

**Figure 1:**
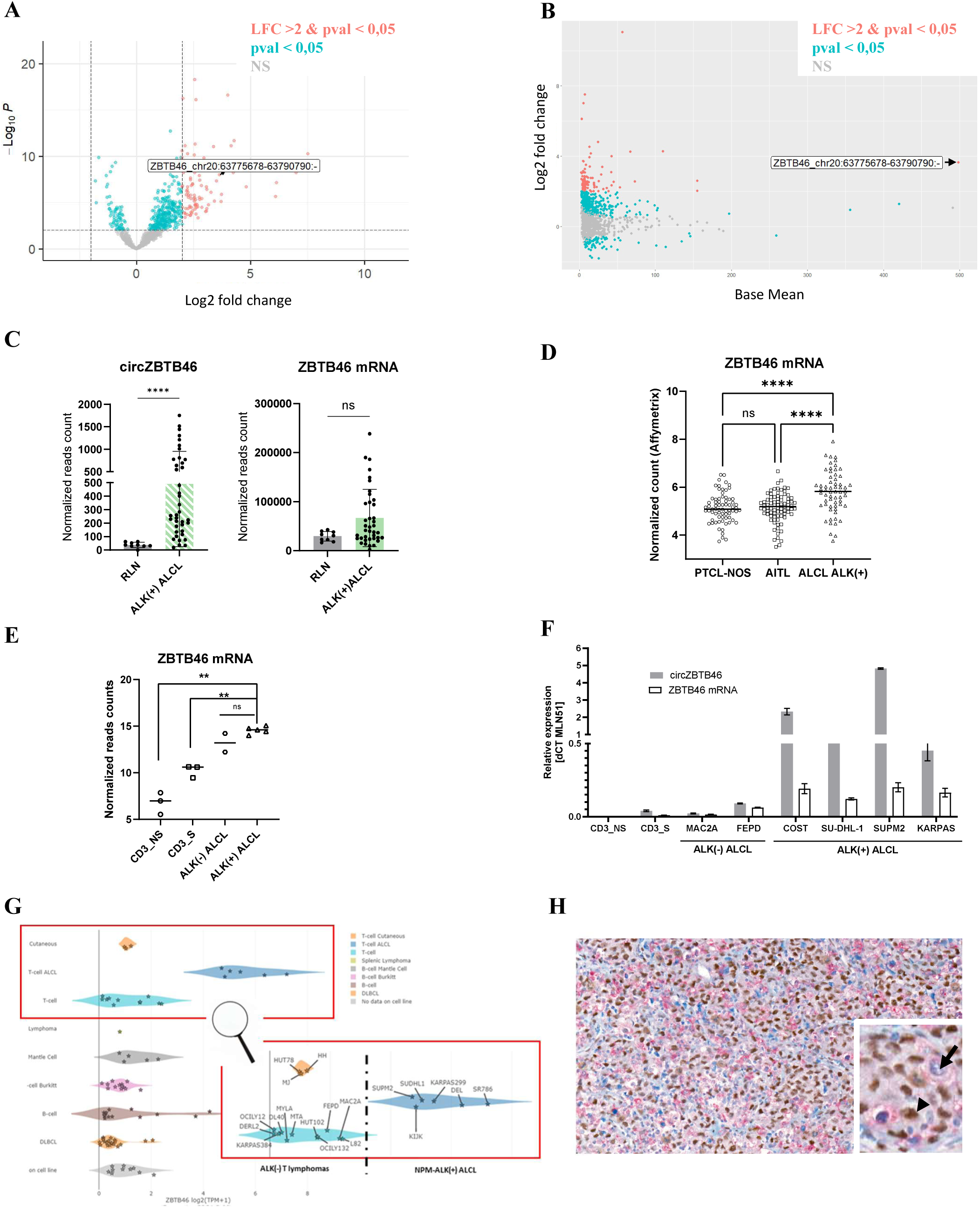
Expression of circZBTB46 and ZBTB46 mRNAs in ALK(+) ALCL primary tumors and normal tissues. **(A)** Volcano plot and **(B)** MA plot of circular RNA expression comparing ALK(+) ALCL primary biopsies (n=39) and healthy tissues (reactive lymph nodes, RLN, n=9). **(C)** expression of circZBTB46 (left) and ZBTB46 mRNA (right) assessed by RNA-Seq in ALK(+) ALCL primary samples versus RLN. **(D)** ZBTB46 mRNA expression (RMA) from microarray datasets in ALK(+) ALCL (n=61), angioimmunoblastic T-cell lymphoma (AITL, n=83) and peripheral T-cell lymphoma not otherwise specified (PTCL-NOS, n=71) primary biopsies. **(E)** ZBTB46 mRNA expression (log2 of the transcript count per million (lenghScaled TPM) from RNA-Seq data in five ALK(+) ALCL cell lines (KARPAS-299, SU-DHL-1, SUP-M2, Pio, COST), two ALK(-) ALCL cell lines (FEPD, MAC-2A) and CD3(+) lymphocytes stimulated (S, n=3) or not (NS, n=3). **(F)** Quantitative real-time PCR (RTLqPCR) analysis of circZBTB46 and ZBTB46 mRNAs in the same cell types. MLN51 was used as an internal control. Values are expressed as 2^(–Δ)Ct relative ratios. Experiments were performed at least in triplicate. Statistical significance was assessed via an unpaired two-tailed Student’s t test with Welch’s correction: *P* < 0.01 (**\*\***), *P* < 0.001 **(\*\*\***), *P* < 0.0001 **(\*\*\*\***), ns = not significant. Data are expressed as means ± SD. **(G)** ZBTB46 mRNA expression across hematological malignancies in the Cancer Cell Line Encyclopedia (CCLE) dataset. **(H)** Representative immunohistochemical image of an ALK(+) ALCL primary tumor showing ZBTB46 (brown, arrowhead) and CD68 (red, arrow) expression. Cell nuclei were counterstained with hematoxylin (blue). Original magnification, ×24.6.

In a second step, published microarray data from peripheral T-cell lymphomas (PTCL)^17^ were analyzed. *ZBTB46* mRNA levels were compared in samples from ALCL (n=61), PTCL-not-otherwise specified (PTCL-NOS, n=71), and angioimmunoblastic T-cell lymphoma (AITL, n=83)^21,22^. The linear *ZBTB46* transcript was found to be significantly upregulated in ALCL compared with other PTCL subtypes (**Fig. 1D**).

Next, expression levels of *ZBTB46* were analyzed in four ALK(+) ALCL cell lines (COST, SU-DHL-1, SUPM2 and KARPAS-299), two ALK(-) cell lines (FEPD and Mac2a) as well as in stimulated (S) and non-stimulated (NS) normal CD3^+^ lymphocytes (n=3 donors). Both RNAseq (**Fig. 1E**) and RTLqPCR (**Fig. 1F**) analyses consistently revealed a strong upregulation of *ZBTB46* mRNA in all ALK(+) ALCL cell lines compared with ALK(-) cell lines and normal CD3+ lymphocytes, independently of stimulation. In addition, RT-qPCR analysis showed a marked overexpression of circZBTB46 in ALK(+) ALCL cell lines compared with controls (**Fig. 1F**). These findings were confirmed at the protein level via Western blotting (**Supplementary Fig. 1**).

Using the Cancer Cell Line Encyclopedia (CCLE) database and the DepMap portal website^23^ we found again that expression of the linear *ZBTB46* transcript was far the highest in ALK(+) ALCL cell lines among lymphomas cell lines (**Fig. 1G**).

Finally, immunohistochemistry on primary biopsies from 19 ALK(+) and 3 ALK(-) ALCL confirmed that ZBTB46 protein expression was restricted to ALK(+) lymphoma cells (**Fig. 1H**), and that no ZBTB46 expression was detected in CD68^+^ cells (macrophages and dendritic cells) (**Fig. 1H**),

Collectively, these results demonstrated that both circZBTB46 and its linear counterpart are aberrantly and quasi-exclusively expressed in ALK(+) ALCL tumoral cells.

### Basic information and characteristics of circZBTB46 in ALK(+) lymphoma cells

As previously reported^18^, circZBTB46 is a 1255-nucleotide-long circRNA produced by backsplicing exons 2 and 3 of the linear *ZBTB46* transcript (ENST00000245663.9, **Fig. 2A**). CircZBTB46 is listed in circBase and circBank as hsa_circ_0002805, and mapped to the genomic coordinates chr20:62407030-62422143 (hg19). Moreover, it is annotated as circZBTB46(2,3).1 according to the standardized nomenclature for eukaryotic circular RNAs^24^. The circular nature of circZBTB46 in the ALK(+) ALCL cell line SU-DHL-1, was confirmed by resistance to exonuclease digestion (RNase R), unlike linear *ZBTB46* mRNA (**Fig. 2B**). Furthermore, upon transcription arrest by actinomycin D, circZBTB46 remained stable for 8h while its linear counterpart declined rapidly (**Fig. 2C**). Finally, subcellular fractionation revealed that circZBTB46 was predominantly localized in the cytoplasm (**Fig. 2D**).

**Figure 2:**
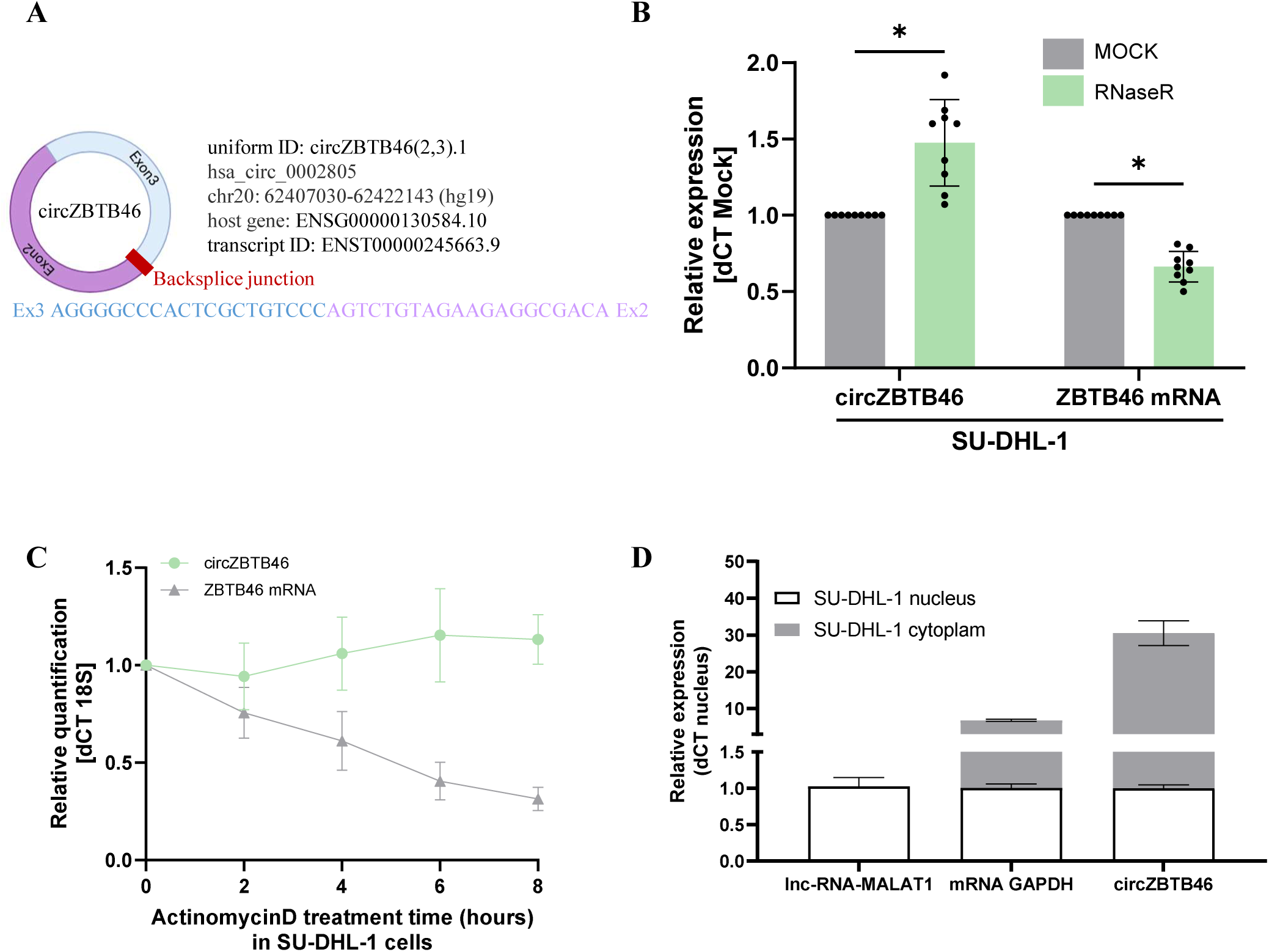
Characterization of circZBTB46 in the SU-DHL-1 ALK(+) ALCL cell line. **(A)** Schematic representation of the circZBTB46 structure with Sanger sequencing of the back-splice junction. **(B)** Quantitative real-time PCR (RTLqPCR) analysis of circZBTB46 and ZBTB46 mRNA expression in cells treated or not with RNase R, a 3′-5′ exonuclease that degrades linear RNAs but spares circular RNAs. **(C)** RTLqPCR analysis after treatment with actinomycin D, an inhibitor of RNA synthesis, to assess transcript stability. 18S rRNA was used as an internal control. Expression values are displayed as 2-ΔΔCt relative ratios. **(D)** Subcellular localization of circZBTB46 and ZBTB46 mRNAs determined by cellular fractionation. MALAT1 and GAPDH mRNAs were used as nuclear and cytoplasmic controls, respectively. All experiments were performed in triplicate. Statistical significance was assessed via an unpaired two-tailed Student t test with Welch correction: *P* < 0.05 (**\***). Data are expressed as means ± SD.

Collectively, these results support that circZBTB46 is a bona fide circular RNA transcript with a cytoplasmic localization.

### The NPM1-ALK/STAT3 axis is responsible for the aberrant accumulation of circRNA and linear *ZBTB46* transcripts in ALK(+) lymphoma cells

The consistent upregulation of both circRNA and linear *ZBTB46* transcripts in NPM1-ALK(+) cell lines and patient primary biopsies suggests that the NPM1-ALK fusion protein plays a central role in driving this overexpression. This was investigated by analyzing *ZBTB46* linear mRNA levels in T-cells ectopically expressing NPM1-ALK, using two complementary models, i) primary human CD4^+^ T-lymphocytes transduced with a lentiviral vector encoding NPM1-ALK (M1 model)^25^ and ii) T-cells engineered to carry the canonical t(2;5)(p23;q35) translocation via CRISPR-Cas9 genome editing (ALKIma1 model)^18^. In both models, previously generated RNAseq datasets were used to demonstrate robust induction of *ZBTB46* mRNA following immortalization and transformation of CD4^+^ T-cells by the NPM1-ALK oncogene (Supplementary **Supplementary Fig. 2A and 2B**). To further determine whether the tyrosine kinase activity of NPM1-ALK is required for this regulation, the ALK(+) ALCL cell line COST was treated with crizotinib. TKI treatment indeed efficiently inhibited ALK kinase activity, as indicated by the loss of NPM1-ALK autophosphorylation (p-ALK). A concomitant reduction in STAT3 activation was evidenced by decreased p-STAT3 levels in Western blot (**Fig. 3A**). RTLqPCR and Western blot further demonstrated a significant downregulation of both circZBTB46 and linear *ZBTB46* transcripts (**Fig. 3B**), as well as reduced ZBTB46 protein levels. (**Fig. 3C**). siRNAs against *NPM1-ALK* (siALK) or *STAT3* (siSTAT3) were then used in two ALK(+) ALCL cell lines, COST and SUP-M2. Efficient knockdown of NPM1-ALK and STAT3 was confirmed at the protein level (**Fig. 3D**). RTLqPCR analysis demonstrated that knockdown of either *NPM1-ALK* or *STAT3* strongly decreased circZBTB46 and linear *ZBTB46* expression in both cell lines (**Fig. 3E**), indicating that NPM1-ALK-dependent STAT3 signaling is critical for driving *ZBTB46* accumulation and circZBTB46 production. Whether STAT3 binds directly to the *ZBTB46* locus was investigated by analyzing publicly available STAT3 ChIP-seq datasets (GSE117164) generated from two ALK(+) ALCL cell lines (JB6 and SU-DHL-1) exposed or not to crizotinib^26^. As a positive control, strong STAT3 binding peaks were observed at the regulatory regions of *IRF4*, a known direct STAT3 target in ALK(+) ALCL cells^27^. Two main STAT3 binding peaks were detected in the *ZBTB46* locus in untreated cells, whereas TKI treatment significantly reduced STAT3 occupancy at these sites, consistent with the loss of STAT3 activation (**Supplementary Fig. 2C**). ChIPLqPCR, using a STAT3-specific antibody in the ALCL cell lines SUP-M2 and COST, and two independent primer pairs within the *ZBTB46* locus, allowed for precise quantifications of STAT3 binding (ZBTB46_1 and ZBTB46_2, **Fig. 3F**). Enrichment of STAT3 binding at its two binding sites was confirmed in the *ZBTB46* locus (**Fig. 3G**).

**Figure 3.**
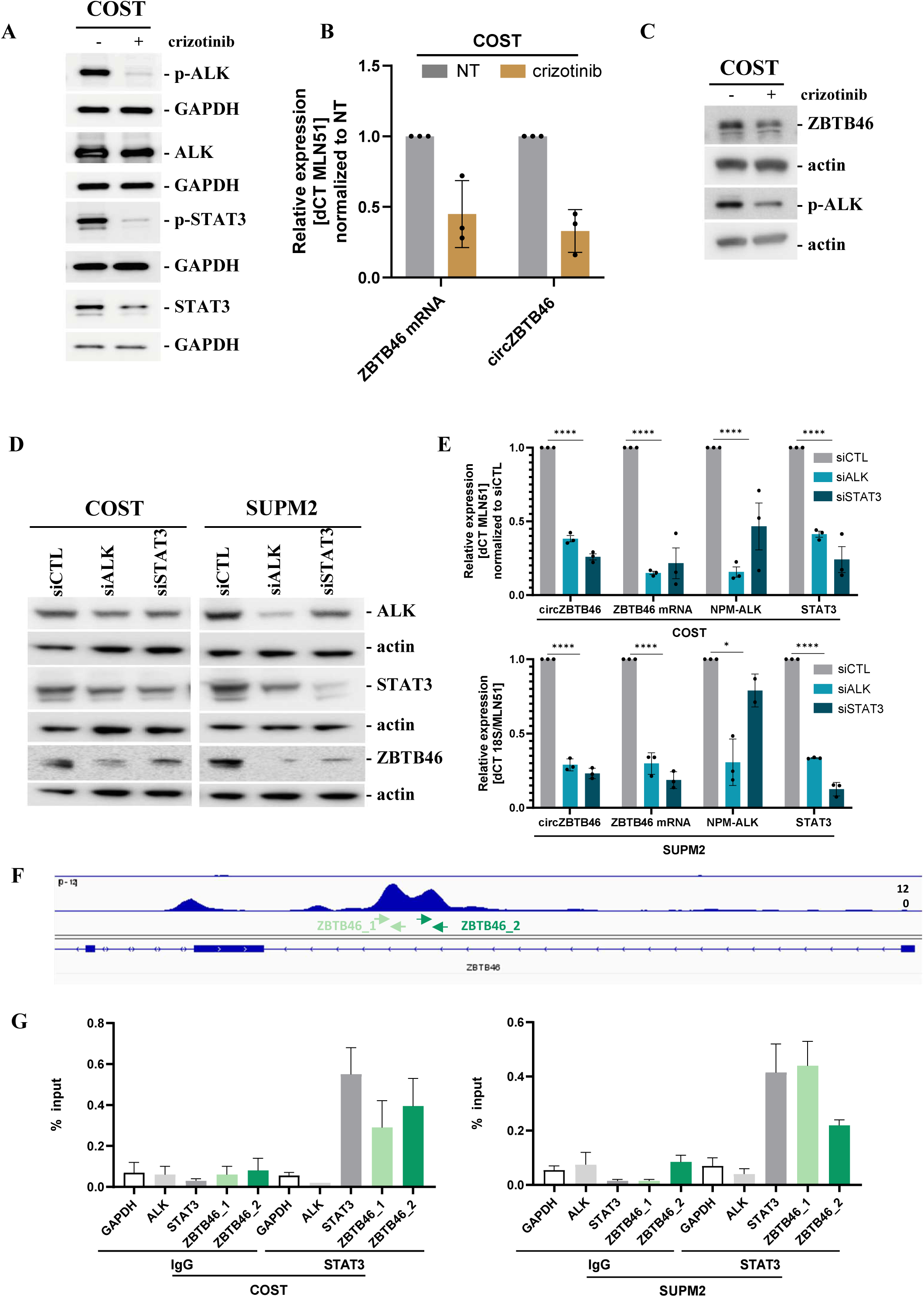
Regulation of *circZBTB46* and *ZBTB46* expression by ALK and STAT3 signaling in ALK(+) ALCL. (**A**) Western blot analysis of total and phosphorylated ALK (ALK, p-ALK) and STAT3 (STAT3, p-STAT3) protein levels in COST cells treated for 48 h with crizotinib (400 μM). (**B**) RTLqPCR analysis of *circZBTB46* and *ZBTB46* mRNA levels following crizotinib treatment (Crizo). (**C**) Western blot analysis of ZBTB46 and p-ALK protein expression under the same conditions. (**D**) Western blot and (**E**) RTLqPCR analysis of ZBTB46, ALK, and STAT3 expression in COST and SUP-M2 cells transfected for 48 h with control siRNA (siCTL), ALK-targeting siRNA (siALK), or STAT3-targeting siRNA (siSTAT3). GAPDH or actin were used as a loading controls. MLN51 served as an internal control for RTLqPCR. mRNA expression values are shown as 2-ΔΔCt relative ratios. Experiments were performed at least in triplicate. (**F**) STAT3 ChIP-seq data from the JB6 cell line showing enrichment of STAT3 at the *ZBTB46* locus (data from GSE117164; ^26^). (**G**) ChIPLqPCR analysis using two primer pairs (ZBTB46_1 and ZBTB46_2) showing STAT3 enrichment at the *ZBTB46* promoter in SUP-M2 and COST cells, represented as % input. STAT3 was a positive control and ALK and GAPDH were used as negative controls. IgG was used as a nonspecific binding control. Experiments were performed at least in triplicate. Representative Western blots from three independent experiments are shown. Statistical significance was assessed via an unpaired two-tailed Student t test with Welch correction: P < 0.05 (*), P < 0.0001 (****). Data are expressed as means ± SD.

Together, these data support a direct regulatory role of the NPM1-ALK/STAT3 axis in the aberrant accumulation of linear and circRNA *ZBTB46* transcripts in ALK (+) ALCL cells.

### CircZBTB46 promotes crizotinib resistance in ALK(+) lymphoma cells

To dissect the respective roles of circZBTB46 and its linear mRNA counterpart, CRISPR-Cas9 knockout models were generated in the COST cell line. Since both the zinc finger domain (ZNF) and nuclear localization signal (NLS) of ZBTB46 are encoded by exon 4, two guide RNAs targeting exons 4 and 5 were used, to induce loss of function of the ZBTB46 protein (**Fig. 4A**). Deletion of exon 4 in three independent clones, Del4-5#C2, #C5 and #E2, was confirmed via genomic PCR (supplementary **Supplementary Fig. 3A**) and RTLqPCR (**Fig. 4B**). This strategy resulted truncated ZBTB46 protein expression, as confirmed in Western blot, using an antibody directed against an epitope encoded by exon 2 (supplementary **Supplementary Fig. 3B**). Additional clones lacking both the functional protein (i.e. mRNA) and circZBTB46 were generated using guide RNAs targeting exons 2 and 5 (**Fig. 4C**), resulting in the deletion of exons 2 to 4. DNA PCR and RTLqPCR confirmed successful deletion in three clones, Del2-5#A2, #A3, and #E2 (**Fig. 4D** and supplementary **Supplementary Fig. 3A**). As expected, these clones exhibited loss of both ZBTB46 protein (supplementary **Supplementary Fig. 3B**) and circRNA (**Fig. 4D**). In immunofluorescence, the full-length ZBTB46 transcription factor was localized in the nucleus of parental wild-type COST cells (**Fig. 4E**), while it was in the cytoplasm of Del4-5 clones (Supplementary **Supplementary Fig. 3C**) and absent in Del2-5 clones (**Fig. 4E** and Supplementary **Supplementary Fig. 3B-C**). The proliferation and survival rates of parental Del4-5 and Del2-5 clones were comparable (Supplementary **Supplementary Fig. 3D-E**). However, after 48h of treatment with crizotinib, both Del4-5 and Del2-5 clones exhibited increased survival compared with parental wild-type cells (**Fig. 4F**). Importantly, Del2-5 clones, lacking circZBTB46 expression, presented significantly less TKI resistance than clones retaining this circRNA (Del4-5). To gain deeper insight into this observation, selective siRNA downregulation of circZBTB46 in the COST and KARPAS-299 cell-lines, was designed to target the back-splice junction and specifically deplete circZBTB46 without affecting the linear *ZBTB46* transcript (siCircZBTB46#1 and #2; **Fig. 5A**). After 48h of crizotinib treatment, both cell lines transfected with circZBTB46-targeting siRNAs exhibited reduced survival compared with control siRNA-treated cells (**Fig. 5B**), supporting the involvement of circZBTB46 in crizotinib resistance in ALK(+) ALCL.

**Figure 4:**
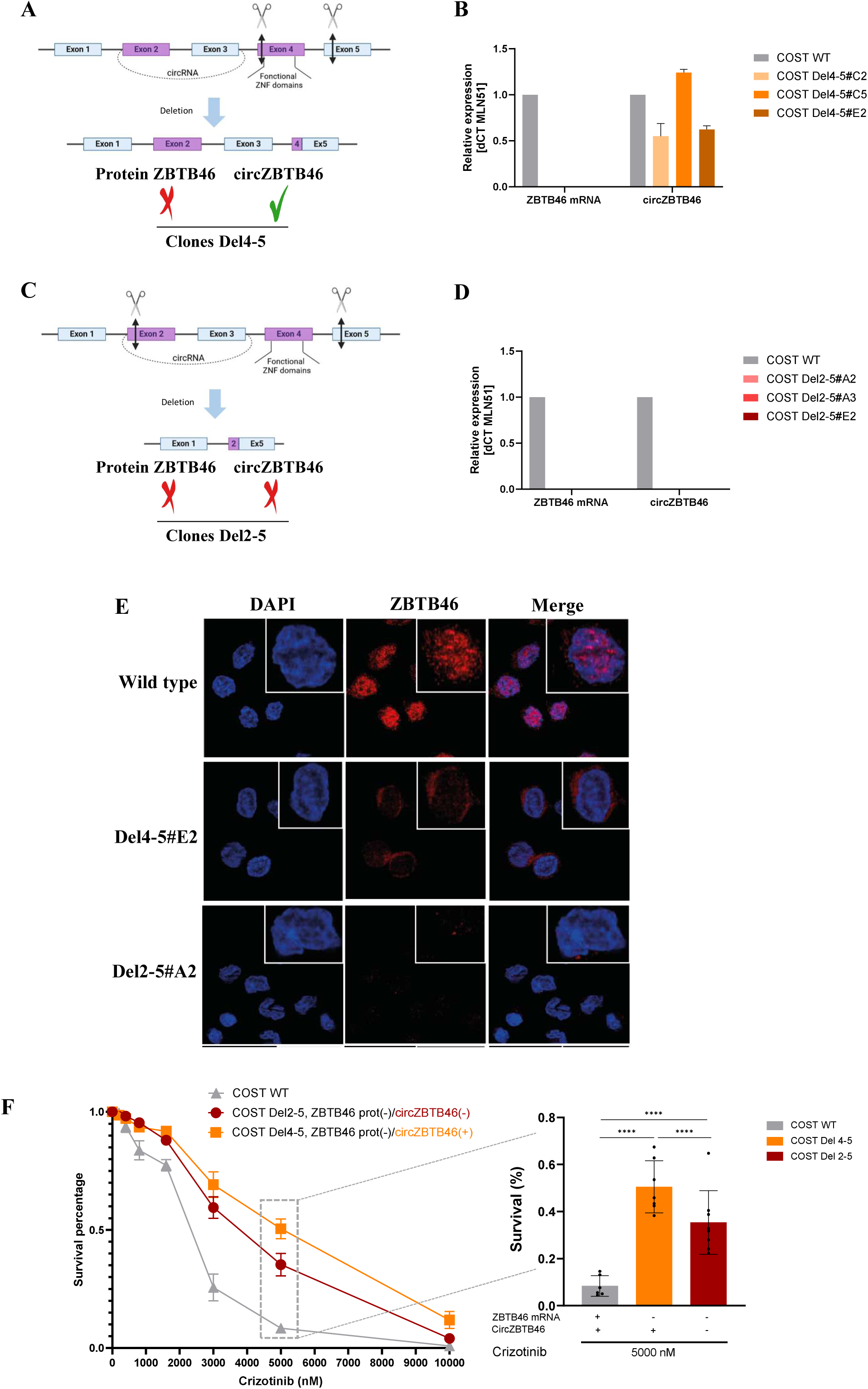
Generation and characterization of ZBTB46 CRISPR/Cas9 knockout models. **(A)** Schematic representation of the genomic deletion between exons 4 and 5 (Del4–5). **(B)** Relative expression levels of *circZBTB46* and *ZBTB46* mRNAs determined by RTLqPCR in COST Del4–5 clones. **(C)** Schematic representation of the genomic deletion between exons 2 and 5 (Del2–5). **(D)** Relative expression levels of *circZBTB46* and *ZBTB46* mRNAs determined by RTLqPCR in COST Del2–5 clones. *MLN51* was used as an internal control for RTLqPCR. Data are expressed as 2-ΔΔCt relative values. All experiments were performed with at least three biological replicates. (**E**) Immunofluorescence analysis of ZBTB46 protein expression in parental (wild-type) COST cells and Del4–5#A2 and Del2–5#E2 clones. Nuclei were counterstained with DAPI (blue). Original magnification, 63×. **(F)** Cell viability was assessed by Annexin V-Pacific blue/PI staining (flow cytometry) in parental COST cells and Del4-5 and Del2-5 clones treated with increasing concentrations of crizotinib (1, 2, 3, 5 and 10 µM) for 48 h.

**Figure 5.**
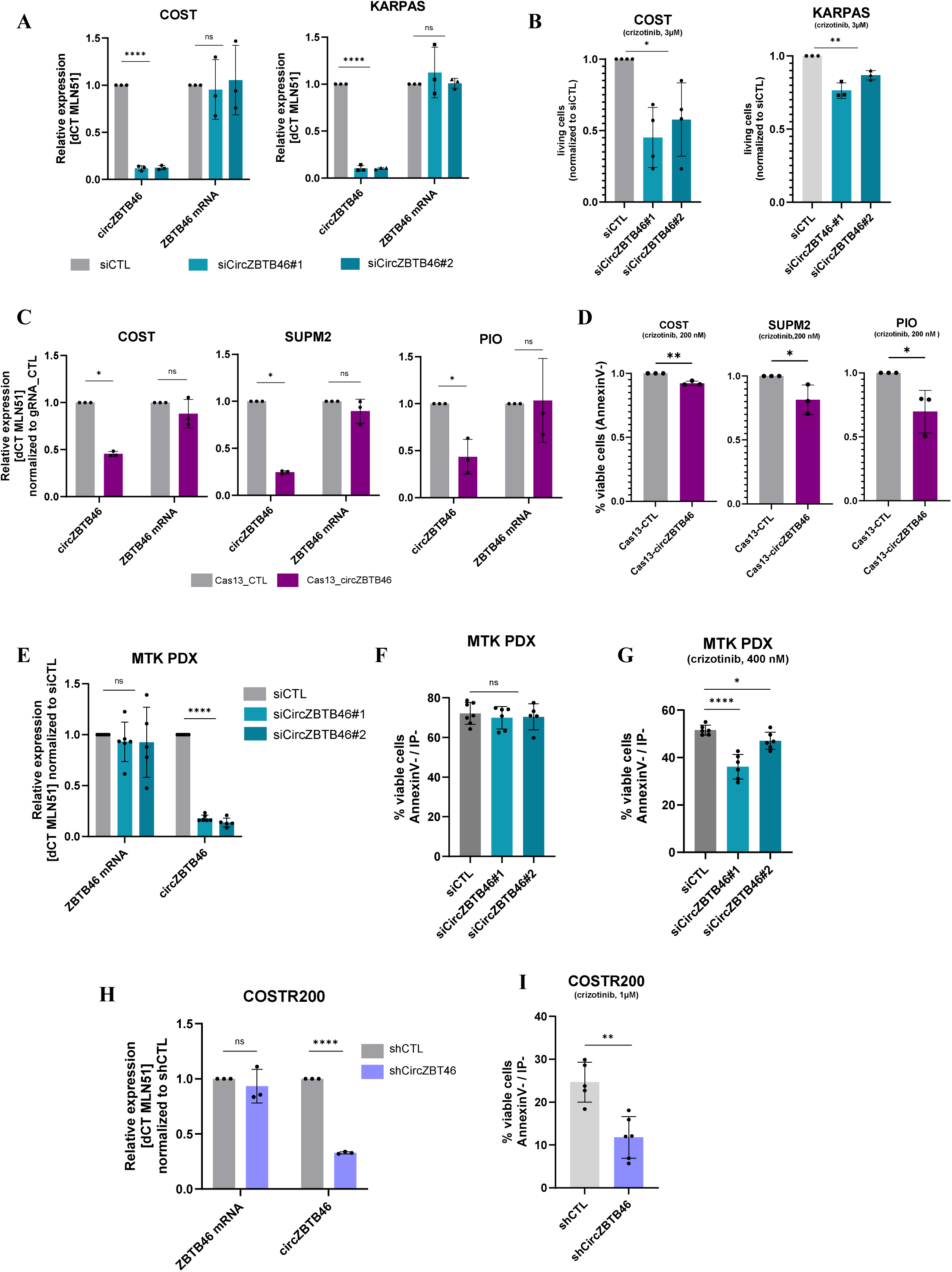
Functional effects of circZBTB46 depletion and sensitivity to crizotinib. **(A)** Relative expression of circZBTB46 and ZBTB46 mRNAs in COST and KARPAS-299 cells transfected for 48 h with control siRNA (siCTL) or two different circZBTB46-targeting siRNAs (siCircZBTB46#1 and #2), as measured by RTLqPCR. **(B)** Viability (Annexin V-Pacific Blue/PI flow cytometry) of COST and KARPAS-299 cells transfected with siCircZBTB46#1 or #2 and subsequently treated with crizotinib (3 µM, 48 h). **(C)** Relative expression of circZBTB46 and ZBTB46 mRNAs in COST, SUPM2 and Pio cells transduced with Cas13 and either control gRNA (CTL) or gRNA targeting circZBTB46, as assessed by RTLqPCR. **(D)** Viability (Annexin V-Pacific Blue/PI flow cytometry) of COST, SUPM2 and Pio cells transduced as described in **(C)** and treated with crizotinib (100 or 200 nM) for 7 days. (**E**) Relative expression of circZBTB46 and ZBTB46 mRNAs in crizotinib-resistant cells established from MTK PDX-derived cells transfected for 48 h with control siRNA (siCTL) or two different siRNAs targeting circZBTB46 (siCircZBTB46#1 and #2), as measured by RTLqPCR. (**F-G**) Cell viability assessed by Annexin V-Pacific Blue/PI staining (flow cytometry) in crizotinib-resistant cells established from MTK PDX-derived cells transfected with siCircZBTB46#1 or #2, without treatment (**F**) or after crizotinib exposure (**G**) (400 nM, 48 h). (**H**) Relative expression of circZBTB46 and ZBTB46 mRNAs in COST crizotinib-resistant cells (COSTR200) after lentiviral transduction with circZBTB46-targeting shRNAs, as measured by RTLqPCR. (**I**) Viability of COSTR200 cells determined by Annexin V/PI staining (flow cytometry) after transduction with circZBTB46 shRNA and subsequent treatment with crizotinib (1,000 nM, 7 days). MLN51 was used as an internal control for RTLqPCR. Expression data are displayed as 2-ΔΔCt relative values. Statistical significance was assessed via an unpaired two-tailed Student t test with Welch correction: *P* < 0.05 (**\***), *P* < 0.01 **(\*\***), *P* < 0.0001 (**\*\*\*\***), ns = not significant. Data are expressed as means ± SEMs.

Based on these results, a CRISPR/Cas13 lentiviral system was used to stably suppress the expression of circZBTB46 (Cas13_circZBTB46) in the COST, SUPM2 and Pio cell lines. This approach resulted in an approximately 50% reduction in circZBTB46 levels without affecting *ZBTB46* mRNA or protein expression (**Fig. 5C**, Supplementary **Supplementary Fig. 4**). Stable circZBTB46 knockdowns were used to assess the effect of prolonged crizotinib exposure (7 days, 100 or 200 nM). Compared with controls (Cas13_CTL), the three transduced cell lines (Cas13_circZBTB46) exhibited significantly reduced viability upon treatment (**Fig. 5D**), indicating increased sensitivity to TKI.

### Loss of circZBTB46 restores the sensitivity of resistant ALK(+) ALCL cells to crizotinib

To explore the therapeutic potential of targeting circZBTB46 in crizotinib-resistant ALK(+) ALCL, siRNA-mediated inactivation of circZBTB46 was performed in MTK PDX-derived cells, obtained from a patient with crizotinib-refractory disease^28^. Two independent siRNAs targeting circZBTB46 were used to ensure specificity of the observed effects (**Fig. 5E**). Silencing circZBTB46 did not affect cell viability in the absence of treatment (**Fig. 5F**). However, after 48h of crizotinib exposure, transfected cells showed increased sensitivity to crizotinib compared with those treated with control siRNAs (**Fig. 5G**). We next generate stable circZBTB46-knockdown cell lines via shRNA lentiviral transduction (shCircZBTB46). Owing to the low transduction efficiency of MTK PDX-derived cells, a previously established crizotinib-resistant derivative of the COST cell line (COSTR200), was used^29^. The effective and specific downregulation of circZBTB46 was confirmed (**Fig. 5H**). As expected, circZBTB46 knockdown sensitized COSTR200 cells to crizotinib *in vitro* (**Fig. 5I**).

### CircZBTB46 induces crizotinib resistance by modulating PIP5K1C gene expression in ALK(+) ALCL

To identify molecular pathways regulated by circZBTB46, RNA-Seq was performed after the transfection of two independent siRNAs targeting circZBTB46 in crizotinib-resistant MTK PDX-derived cells. Both siRNAs induced comparable transcriptional changes, particularly downregulation of PIP5K1C and DHRS9 (|logLFold Change| ≥ 0.5, P < 0.05) (**Fig. 6A**). The downregulation of PIP5K1C was validated by RTLqPCR and Western blotting, confirming reduced expression at both mRNA and protein levels in all cellular models previously described (**Fig. 6B-D**). By contrast, DHRS9 downregulation was not consistently observed (data not shown).

**Figure 6.**
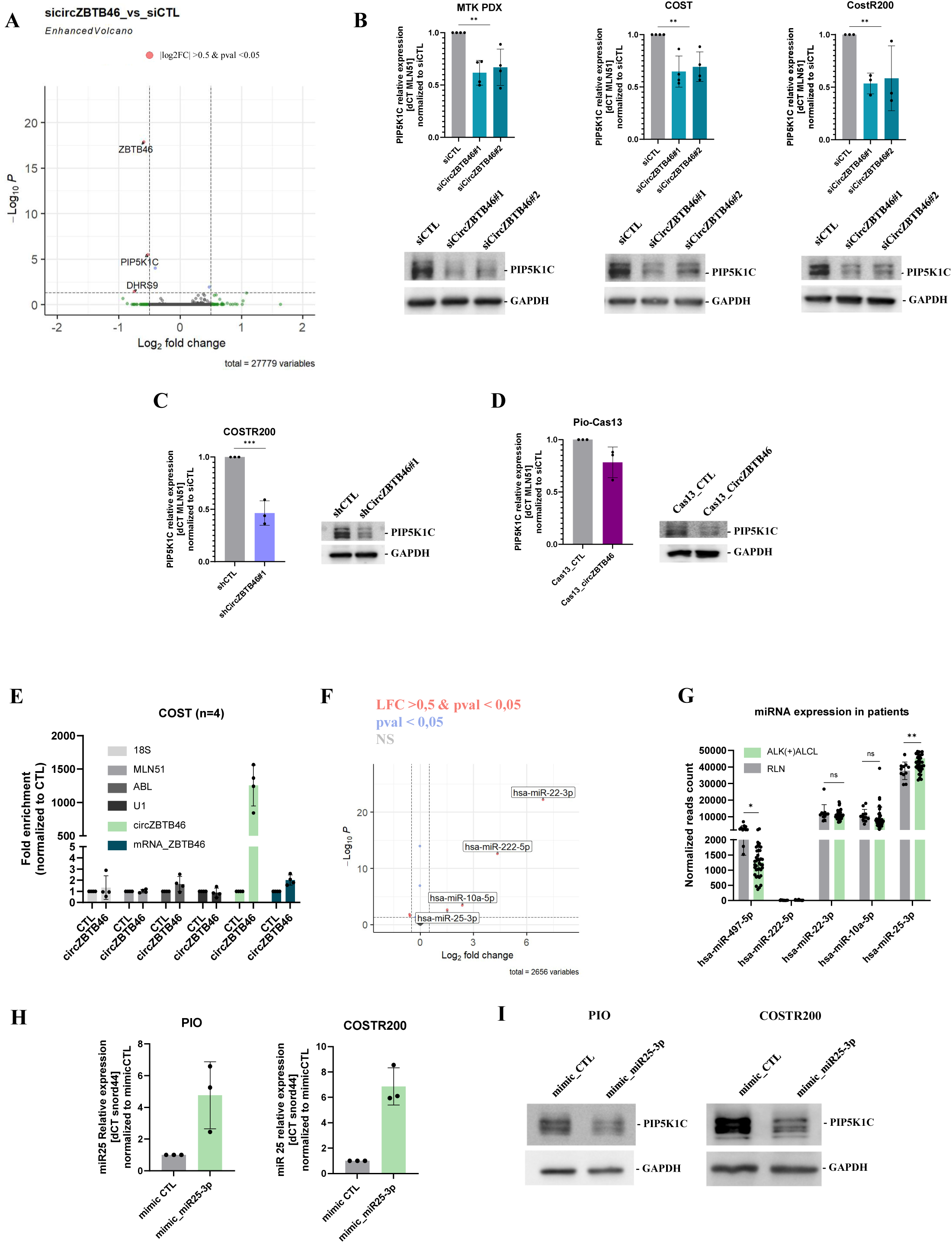
circZBTB46 modulates PIP5K1C by sponging miR-25. **(A)** Volcano plot showing differentially expressed genes in crizotinib-resistant cells established from MTK PDX-derived cells transfected with control siRNA (siCTL) or siRNAs targeting circZBTB46 (siCircZBTB46#1 and #2). **(B)** Expression of PIP5K1C at the mRNA (top panel, RTLqPCR) and protein (bottom panel, Western blot) levels in COST, MTK-PDX and COSTR200 ALK(+) ALCL cells after circZBTB46 siRNA transfection (48 h). **(C)** Expression of PIP5K1C at the mRNA (left panel) and protein (right panel) levels in stable COSTR200 cells expressing either control shRNA (CTL) or shRNA targeting circZBTB46, as assessed by RTLqPCR and Western blot, respectively. **(D)** Expression of PIP5K1C in Pio ALK(+) ALCL cells stably expressing Cas13 with control gRNA (CTL) or gRNA targeting circZBTB46, as determined by RTLqPCR (left panel) and immunoblotting (right panel), respectively. **(E)** RNA enrichment after RNA pull-down using biotinylated probes targeting circZBTB46 or control probes**. (F)** Volcano plot showing in the COST cell line differentially enriched RNAs in circZBTB46 pull-down compared to control, highlighting specific microRNAs enriched in the circZBTB46 fraction. **(G)** Relative expression levels of microRNAs (miR_497-5p, miR_222-5p, miR_22-3p, miR_10a-5p, and miR_25-3p) in ALK(+) ALCL primary samples versus reactive lymph nodes (RLNs), as assessed by small RNA sequencing. **(H)** Relative expression of miR25-3p in crizotinib-sensitive (Pio) and resistant COSTR200 cells transfected with a negative control mimic (mimicCTL) or mimic_25-3p measured by RT-qPCR. Snord44 was used as an internal control. Data are shown as 2-ΔΔCt relative values**. (I)** Western blot analysis of PIP5K1C protein levels in PIO and COSTR200 cells transfected with a miR_25-3p mimic or control mimic. GAPDH served as a loading control. Experiments were performed in triplicate. Data are presented as mean ± SEM. Statistical significance was determined using an unpaired two-tailed Student t test with Welch correction: *P* < 0.05 (**\***), *P* < 0.001 (**\*\***), ns = not significant

Contribution of PIP5K1C to crizotinib resistance in ALK(+) ALCL was explored by CRISPR interference (CRISPRi) technology. COSTR200 cell line were transduced with dCas9-KRAB-MeCP2 fusion proteins using guide RNAs targeting the PIP5K1C promoter to achieve transcriptional silencing. As shown in **Supplementary Fig. 5A**, gRNA-mediated silencing of PIP5K1C resulted in a significant reduction in PIP5K1C protein expression. Compared with control cells, PIP5K1C-silenced cells exhibited a slight decreased viability *in vitro* (**Supplementary Fig. 5B**). These results indicate that regulation of PIP5K1C by circZBTB46 participates in promoting resistance to ALK inhibition in ALK(+) ALCL.

### CircZBTB46 regulates PIP5K1C by sponging miR25-3 in ALK(+) ALCL cells

After RNA pull-down was performed using a back-splice junction-specific probe in ALK(+) ALCL cells, RTLqPCR (**Fig. 6E**) confirmed a successful enrichment of circZBTB46. Mass spectrometry analysis of associated proteins did not reveal any specific interactors (data not shown), whereas small RNA sequencing identified four miRNAs (cutoff of |log2Fold Change|≥ 2 and a *P* value<0.05), namely, hsa-miR-22-3p, hsa-miR-222-5p, hsa-miR-10a-5p, and hsa-miR-25-3p, that were significantly enriched in the circZBTB46 pull-down compared with control conditions (**Fig. 6F**). With the exception of hsa_miR_222-5p, the other three miRNAs were expressed in ALK(+) ALCL patients (**Fig. 6G**), hsa-miR-25-3p being notably overexpressed compared with RLN. Using miRNA–mRNA interaction prediction algorithms (miRDB, TargetScanHuman and miRTargetLink 2.0), PIP5K1C was identified as a putative target of miR-25-3p (**Supplementary Fig. 5C-D**). Consistently, transfection of a miR-25-3p mimic in crizotinib-sensitive (Pio) and crizotinib-resistant (COSTR200) cells, as validated by RTLqPCR analysis (**Fig. 6H**), led to a reduction in PIP5K1C protein level (**Fig. 6I**). Conversely, transfection of a miR-25-3p inhibitor significantly increased *PIP5K1C* mRNA and protein levels as observed in in COSTR200 cells, further validating this regulatory pathway (**Supplementary Fig. 5E-F)**. Together, these data indicate that circZBTB46 modulates PIP5K1C levels via miR-25-3p sequestration, uncovering a novel competing endogenous RNA axis in ALK(+) ALCL.

### Targeting circZBTB46 in ALK(+) ALCL cells reduces tumor growth *in vivo*

*In vitro* experiments revealed that circZBTB46 is involved in crizotinib resistance in ALK(+) ALCL cells. To evaluate its potential as therapeutic target, the effect of circZBTB46 targeting on tumor growth was analyzed after subcutaneous injection in NSG mice.

First, CRIPSR/Cas9 crizotinib-sensitive models were engrafted. Del2-5 and Del4-5 clones formed tumors at comparable rates (**Fig. 7A**). Daily administration of crizotinib, started five days after transplantation, initially reduced the tumor volume in all groups. However, by day 15, all tumors derived from ZBTB46 RNA-depleted clones resumed growth, whereas those derived from parental (WT) cells continued to regress (**Fig. 7B**). Notably, tumors derived from Del2-5 clones lacking circZBTB46 exhibited slower growth than those derived from circZBTB46(+) clones (Del4-5), corroborating *in vitro* findings. Next, Cas13-modified crizotinib-sensitive Pio cells were xenografted into NSG mice and treated daily with crizotinib since nine days after injection. Tumor growth was delayed in mice bearing circZBTB46-deficient tumors (**Fig. 7C**). As PIP5K1C was identified as a downstream target of circZBTB46, PIP5K1C CRISPRi models were also injected in NSG mice. This resulted in a slight reduction of tumor growth upon crizotinib administration (**Supplementary Fig. 6**), highlighting that PIP5K1C as a mediator in TKI-resistance phenotype. Finally, the crizotinib resistant shCircZBTB46 models were injected to determine whether targeting circZBTB46 in crizotinib-resistant cells could restore drug sensitivity. Mice were daily treated with crizotinib or vehicle. In the absence of TKI treatment, tumor growth did not differ between the groups. However, upon crizotinib treatment, tumors derived from shCircZBTB46 cells exhibited significantly reduced growth (**Fig. 7D**). All these results support the contribution of circZBTB46 to crizotinib resistance in ALK(+) ALCL and identify circZBTB46 as an interesting therapeutic target of TKI-resistance to in ALK(+) ALCL disease.

**Figure 7.**
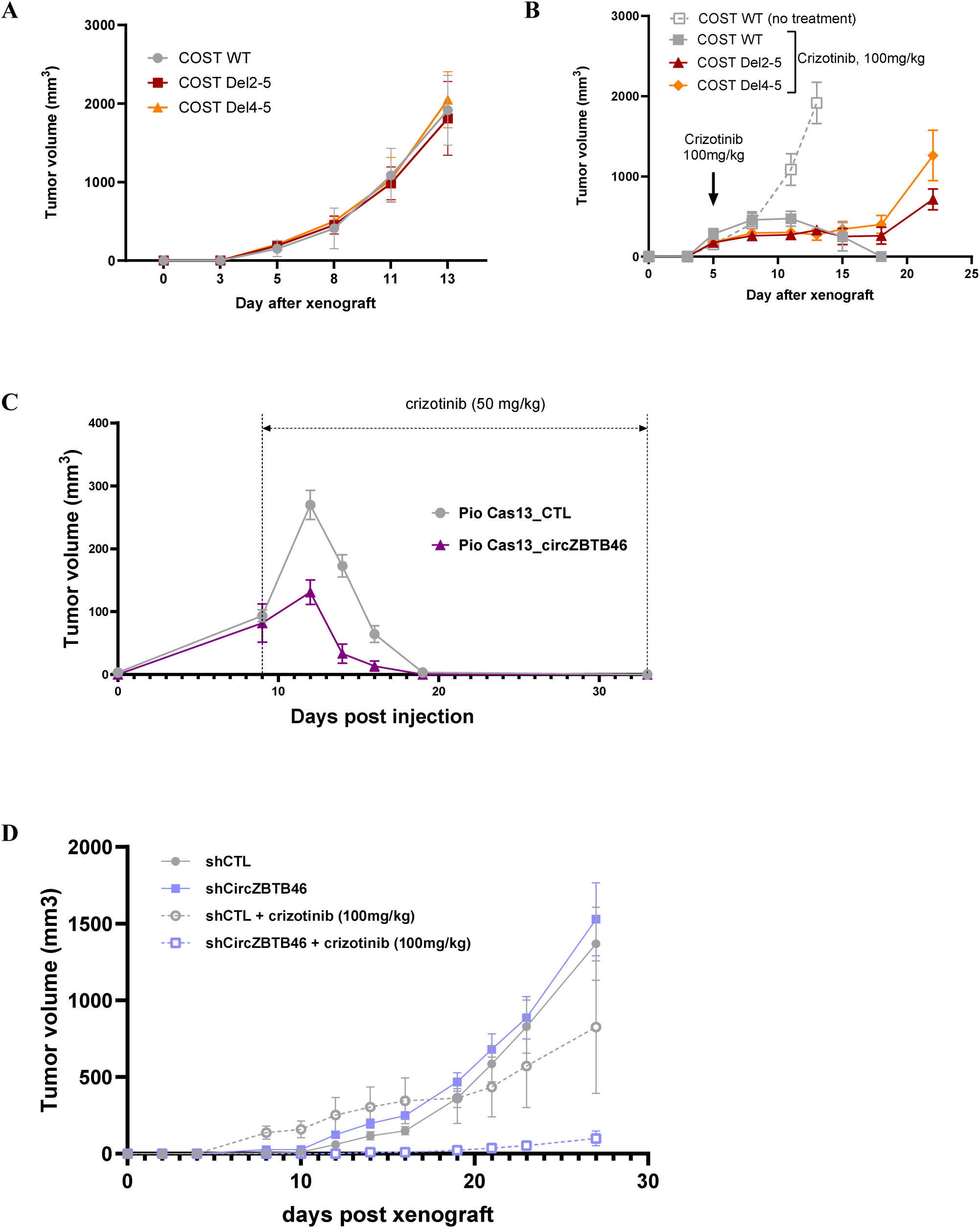
circZBTB46 modulates crizotinib resistance *in vivo* Tumor growth in NSG mice (n=8) injected subcutaneously **(A-B)** with COST cells **(A)** untreated or **(B)** treated with crizotinib (5 mg/kg/day); **C**) with Pio cells expressing Cas13 with either a nontargeting control gRNA (Cas13_CTL) or a gRNA targeting circZBTB46 (Cas13_circZBTB46) and (**D**) with COSTR200 cells transduced with or without circZBTB46 shRNA with or without crizotinib treatment (100 mg/kg/day). Parental cells were included as a control. Tumor volume was monitored over time via calipers and is expressed as the mean ± SD. Statistical significance was assessed via an unpaired two-tailed Student’s t test with Welch correction: *P* < 0.05 (**\***), *P* < 0.01 **(\*\***), *P* < 0.0001 (**\*\*\*\***), ns = not significant. Data are expressed as means ± SD.

## Discussion

This study identifies the circular RNA, hsa_circ_0002805 (circZBTB46), as a key driver of crizotinib resistance in ALK-positive anaplastic large cell lymphoma (ALK(+) ALCL). This circRNA, derived from exons 2 and 3 of the ZBTB46 gene, was found to be significantly and specifically upregulated in this lymphoma type. The upregulation is driven transcriptionally by the STAT3 pathway, which is activated by the oncogenic NPM1-ALK fusion protein. Using a combination of CRISPR/Cas and RNAi technologies, we functionally dissected the role of circZBTB46, finding that it, unlike its linear protein-coding counterpart, promotes resistance to crizotinib, an ALK inhibitor. This was demonstrated in both in vitro and in vivo models. Depleting circZBTB46 in resistant cells restored their sensitivity to crizotinib, highlighting its potential as a therapeutic target. Transcriptomic analysis revealed that PIP5K1C is a downstream effector of the circZBTB46-mediated resistance. Mechanistically, circZBTB46 acts as a molecular sponge for miR-25-3p, a microRNA that normally suppresses PIP5K1C expression. By sequestering miR-25-3p, circZBTB46 de-represses PIP5K1C, promoting survival signals and contributing to drug resistance. This proposed circZBTB46–miR-25-3p–PIP5K1C axis is a novel resistance mechanism in hematologic malignancies.

ZBTB46 is a transcription factor normally restricted to dendritic cell precursors, where it maintains their immature state^20,30,31,34,35^. It is also found in non-proliferative endothelial cells, where it inhibits cell proliferation^32^. Emerging evidence, however, points to a context-dependent role for ZBTB46 in cancer^33–35^. In acute myeloid leukemia (AML), elevated ZBTB46 is associated with poor outcomes^36^, and despite not being essential for normal hematopoiesis, it is crucial for the survival and proliferation of AML cells^37^.

Similarly, circZBTB46 exhibits context-dependent functions across different diseases. In atherosclerosis, circZBTB46 promotes disease progression by activating the AKT/mTOR pathway^38^. In this context, circZBTB46 depletion induces apoptosis and impairs cell migration, highlighting its importance in vascular pathology. In contrast, this study found no effect on AKT/mTOR signaling in ALK(+) ALCL cells upon circZBTB46 silencing (data not shown).

In AML, circZBTB46 has been identified as an oncogenic effector that promotes cell survival. It acts as a competing endogenous RNA, sponging both hsa-miR-326 and hsa-miR-671-5p. This activity increases the expression of stearoyl-CoA desaturase 1 (SCD1), an enzyme that protects cells from lipid peroxidation and ferroptotic cell death.These findings establish circZBTB46 as a novel oncogenic effector in AML and reveal an unrecognized *ZBTB46*-derived noncoding RNA axis that governs cell fate through ferroptosis-related mechanisms^36^.

Notably, both *ZBTB46* linear RNA and circZBTB46 were found to be expressed at markedly higher levels in ALK(+) ALCL than in AML. This was demonstrated by comparative transcriptomic analyses using a k-mer-based approach^39,40^ applied to in-house RNA-seq data from primary biopsy samples, including ALK(+) ALCL and AML samples (IUCT-AML cohort) (**Supplementary Fig. 7**)^41,42^. RLN and established cell lines (AML and ALK(+) ALCL), including *NPM1-ALK*–transformed immortalized T-cell models^18,25^, and healthy T lymphocytes confirmed these data. This unexpected expression pattern underscores the context-dependent transcriptional regulation and functional plasticity of the *ZBTB46* locus across distinct hematological malignancies. In fact, this study expands knowledge on the pathophysiological oncogenic potential of ALK(+) ALCL by demonstrating the role of circZBTB46 in promoting resistance to crizotinib. It was shown here that circZBTB46 is transcriptionally upregulated via the NPM1-ALK/STAT3 signaling axis, thereby linking oncogenic kinase activity to circular RNA-mediated drug resistance. These findings suggest that circZBTB46 acts as a context-specific regulator of therapy resistance by engaging survival pathways, making it a potential therapeutic target in ALK-driven hematological malignancies. In this context, PIP5K1C appears as a downstream target of circZBTB46. miR-25-3p was identified as a candidate intermediary. miR-25-3p has previously been shown to target PIP5K1C in prostate cancer^43^. These findings support a model in which circZBTB46 may act as a molecular sponge for miR-25-3p, thereby derepressing PIP5K1C expression and promoting survival signaling. Thus, circZBTB46 might contribute to crizotinib resistance in ALK(+) ALCL through a miR-25-3p–PIP5K1C axis, a mechanism not previously appreciated in hematologic malignancies.

Interestingly, PIP5K1C was found to play a role in conferring resistance to ALK inhibition in ALCL. This lipid kinase catalyzes the phosphorylation of phosphatidylinositol 4-phosphate (PI4P) to produce phosphatidylinositol 4,5-bisphosphate (PI(4,5)PL), which plays a key role in remodeling the actin cytoskeleton, intracellular trafficking and cell migration^44^. PIP5K1C phosphorylation at serine 448 has been suggested to be a potential biomarker for breast cancer progression, particularly in invasive breast cancer^45^. Compared with their sensitive counterparts, melanoma cells that are resistant to the PIKFYVE inhibitor WX8 exhibit elevated expression of PIP5K1C, and this resistance can be reversed by either siRNA-mediated knockdown of PIP5K1C or pharmacological inhibition of its kinase activity^46^. These findings suggest that PIP5K1C could mediate therapeutic resistance in multiple cancer types including hematological neoplasms. Therefore, focusing on PIP5K1C in crizotinib-resistant ALCL cells could be an effective way to overcome resistance.

## Methods

### Cell line culture

The ALCL cell lines KARPAS-299, SU-DHL-1 SUPM2, Pio and COST, which carry the t(2;5)(p23;q35) translocation, were obtained from the DSMZ (German Collection of Microorganisms and Cell Culture, Leibnitz, Germany) or established locally ^29^. The MTK PDX-derived cell line was provided by J. Mathews and S. Turner ^28^, and the crizotinib-resistant KARPAS-299-CR06 cell line was provided by C. Gambacorti-Passerini ^47^. COST cells were exposed to increasing concentrations of crizotinib, ranging from 25 nM to 200 nM. At each step, the dose was increased only when treated cells achieved a proliferation rate comparable to that of untreated wild-type (WT) cells. All ALK(+) ALCL cell lines and PDXs were cultured in RPMI-1640 medium (Invitrogen, Waltham, MA #61870044) supplemented with 20% FBS (Pan Biotech, Aidenbach, Germany #500105M1M). For crizotinib-resistant cell lines, crizotinib was added to the medium twice a week at a concentration of 200 nM for COSTR200 and 600 nM for KARPAS-299-CR06.

### Crizotinib, siRNA and mimic treatments

Cells were seeded at 2x10^5^ cells/mL and treated at 24h with different doses of crizotinib (Selleckchem, Houston, TX, PF-02341066 #S1068). Aliquotes of 3.10^6^ cells were transfected with 70 to 100 nmol siRNA/mimic (listed in **Supplementary Table S1**) using Amaxa4D nucleofector (Primary Cell kit P3, CA-137 program, Lonza Bioscience, Antwerp, Belgium).

### CRISPR model/shRNA model cell lines

Guide RNAs for CRISPR/Cas13 models were cloned and inserted into the pSLQ5465_pHR_hU6-crScaffold_EF1a-PuroR-T2A-BFP backbone (Addgene, Wartertown, MA, #155307) via BbsI digestion. Guide RNAs for CRISPRi were subsequently cloned and inserted into the pKLV2-U6gRNA5(BbsI)-PGKpuro2ABFP-W backbone (Addgene #67974) via BbsI digestion. shRNAs were cloned and inserted into the pLKO.1 puro backbone (Addgene #8453) via AgeI and EcoRI digestion. All sequences are listed in **Supplementary Table S1**. Lentiviruses were produced in HEK293T cells. The cells were transfected with psPAX2 (Addgene #12260), pMD2. G (Addegene # 12259) and the plasmid of interest using Lipofectamine 2000 and 24 µg total DNA (with a ratio of 1:1:1). Lentiviruses were harvested 48 h after transfection and used directly on ALCL ALK(+) cells. A total of 10^6^ cells were transduced with 1 mL of lentiviral supernatant supplemented with 10 µg/mL polybren (Sigma, St Louis, MO, #TR-1003-G). Cells were spun-inoculated for 2 hours at 1000 rpm and 37°C, after which 1 mL of fresh medium was added. For the CRISPR/Cas13 model, cells were first transduced with pXR001:EF1a-CasRx-2A-EGFP lentivirus (Addgene #109049) and then with pSLQ5465_pHR_hU6-crScaffold_EF1a-PuroR-T2A-BFP lentivirus containing guides of interest. The transduced cells were then selected with puromycin (1 µg/mL). For CRISPRi models, cells were first transduced with dCas9-KRAB-MeCP2 lentivirus (Addgene #110821) and then with pKLV2-U6gRNA5(BbsI)-PGKpuro2ABFP-W lentivirus containing guides of interest. The transduced cells were then selected with puromycin (1 µg/mL). For the shRNA model, cells were transduced with pLKO.1 puro lentivirus containing the shRNA sequence of interest.

### RNA extraction, reverse transcription and quantitative PCR

Total RNA was extracted from 5 to 10x10^6^ cells via TRIzol reagent (Invitrogen # 15596018) followed by a Direct-zol™ RNA Miniprep Plus Kit (Zymo, Orange, CA #R2072) following the manufacturer’s instructions. Typically, 1 µg of total RNA was reverse transcribed via the ProtoScript® II First Strand cDNA Synthesis Kit (NEB, Ipswitch, MA, #E6560L) following manufacturer instructions. cDNA was next diluted with 5 in water, and 2 µL were used for quantitative PCR analysis. qPCR was performed via the Master Mix Select SYBR™ (Applied Biosystems, Waltham, MA, #4472908) in a 10 µL total reaction. The PCR program was as follows: 50°C for 2 min, 95°C for 2 min, and 40 cycles of 95°C for 15 s and 60°C for 1 min. All primer sequences are listed in **Supplementary Table S1.**

### RNA dataset generation and analysis

Microarray data (.CEL files) for patients with ALCL, AITL and PTCL-NOS were retrieved from the translational T-cell lymphoma research consortium (TENOMIC) of the LYmphoma Study Association (LYSA) and PAIR lymphoma project, previously available ^17,25^. Raw data were processed with the *affy* R package and the *rma* function to transform probe intensities into normalized expression values by robust multi-array average (RMA) (Irizarry et al Biostatistics 2003). To get by-gene summarized expression values, the collapse microarray tool was used (https://sites.google.com/site/fredsoftwares/products/collapse-microarray?authuser=0) with the provided Affymetrix U133+2 table corresponding to the chip reference.

The Ribo zero full RNA sequencing for 39 ALCL patient samples and 9 RLN samples was previously published (GSE160123, ^17^). For all other samples, the same library preparation was performed. Pooled library prep samples were subsequently sequenced on a NovaSeq 6000 Illumina (San Diego, CA), corresponding to 1x30 million 100-base reads per sample after demultiplexing. For ZBTB46 mRNA expression analysis shown in Fig. 1E, reads pseudoalignment was performed with Kallisto ^48^ (version 0.46.1) using the v108 Ensembl transcriptome, followed by *tximport* ^49^ for computing gene-level TPM values, normalized by gene length and library size (lengthScaledTPM). De novo identification of circRNAs was performed according to a standard protocol, using the CIRI2 algorithm with the human reference genome GRCh38 ^50^.The circRNA-specific junction reads are non-colinear and are therefore typically found within reads that do not align to the genome. CIRI2 retrieves this “trashed” information to identify and distinguish circular transcripts from linear transcripts ^50^. Differential expression analysis was performed via the DESeq2 tool ^51^.

For all other samples, the reads were aligned with hisat2 ^52^ on the hg38 human genome and expression was quantified via featureCounts ^53^. Differential expression analysis was performed via the DESeq2 tool ^51^.

SmallRNA library preparation was performed according to manufacturer recommendations (QIAseq miRNA Library Kit from Qiagen, LesUlis, France). The final pooled library preparation samples were sequenced on Novaseq 6000 Illumina corresponding to 1x30M 100bases reads per sample after demultiplexing. The quantification of miRNAs was carried out according to recommendations by Potla et al ^54^.

### Chromatin immunoprecipitation

Fifteen million cells were fixed in 2% paraformaldehyde. Chromatin immunoprecipitation was then performed via a ChIP-IT^®^ Express Kit (Active Motif, Shangai, China, #53008) according to manufacturer instructions. Sonication was performed on VibraCell (Sonics & Materials, Newton, CT, #75186) 10 times for 20s at a power of 25s (with a 20s pause in between). Immunoprecipitation was performed using an anti-STAT3 antibody (#12640S, RRID: AB_2629499) and an anti-rabbit IgG antibody (#2729, RRID: AB_1031062) from Cell Signaling Technology (Danvers, MA).

### RNA pulldown

iDRIP was performed as previously described ^55^. Probe sequences are listed in **Extended Data** Table S1.

### Xenograft tumor assay

Mice were housed under pathogen-free conditions in an animal room at constant temperature (20°C-22°C) with a 12-h light/dark cycle and free access to food and water. A total of 3×10^6^ cells were injected subcutaneously into one flank of 5-week-old female NSG mice (Janvier Labs, Le Genest Saint Isle, France). Mouse body weight and tumor volumes were measured 3 times a week with calipers via the following formula: length × width^2^× π/6. At the end of the experiment, the mice (8 per group) were humanely sacrificed. For crizotinib-resistant cells, mice were treated with 100 mg/kg/day crizotinib starting on the day following injection. For crizotinib-sensitive cells, mice were treated with 50 mg/kg/day crizotinib, starting when the tumor size reached 300 mm^3^.

### Statistical analyses

Data are expressed as mean ± standard error of the mean (SEM) of at least three independent experiments. Graphs and statistical analyses were performed via GraphPad Prism 10 (GraphPad Software Inc., La Jolla, CA). P values were calculated by using an unpaired two-tailed Student *T* test with Welch correction.

### CircRNA validation by PCR and Sanger sequencing

The validation of circRNA back-splice junctions, circularity, and stability were performed as previously. In brief, PCR amplification was followed by Sanger sequencing to confirm the back splicing junction and the full circRNA sequence. Circularity and exonuclease resistance were assessed via RNase R treatment, whereas transcript stability was evaluated following exposure to actinomycin D. Relative expression levels were determined via RTLqPCR via the 2-ΔΔCt method.

### Western blot analysis

Whole-cell extracts were prepared with protein lysis buffer (50 mM Tris-HCl pH 7.4, 1%Triton X-100, 0.1% SDS, 150 mM NaCl, 1 mM EDTA, and 1 mM DTT), supplemented with « Complete cocktail protease inhibitor tablets » (Roche, Vienna, Austria, #11697498001) and « Halt Protease and Phosphatase Inhibitor Cocktail » (Thermo Fisher Scientific, Waltham, MA #78440). Typically, 30 μg of protein were loaded on 10% acrylamide gel (w/v) Tris-HCl SDS PAGE. After transfer, membranes were stained with anti-Phospho-ALK (Tyr1604 #3341 RRID: AB_331047), anti-ALK (31F12 #3791, RRID: AB_1950402), anti-P-STAT3 (Tyr705 #9131, RRID: AB_331586), anti-STAT3 (#4904, RRID: AB_331269), anti-PIP5K1C (#3296, RRID:AB_2164719) from Cell Signaling and anti-ZBTB46 (Sigma Aldrich, St Louis, MO, #HPA013997, RRID: AB_1858961, an antibody directed against an epitope encoded by exon 2) from Atlas Antibodies. Anti-Tubulin (Santa Cruz, Dallas, TX, #sc-53029 RRID : AB_793541) Anti-actin (Sigma, #A5316, RRID : AB_476743) and anti-GAPDH (Santa-Cruz, #sc-32233, RRID : AB_627679) antibodies were used as loading controls. Secondary antibodies were HRP-conjugated, and blot developed with ECL prime (BIO-RAD, Hercules, CA, # 1705060).

### Immunohistochemistry

ALK and CD68 protein expression was assessed by immunohistochemistry on formalin-fixed, paraffin-embedded (FFPE) tissue sections. ALK and CD68 expression were detected via immunohistochemistry via a polyclonal rabbit anti-ALK antibody (clone SP8; 1:100 dilution; Lab Vision Corporation, Fremont, CA) and a monoclonal mouse anti-human CD68 antibody (clone PG-M1; Dako, Les Ulis, France), respectively. Signal visualization was achieved via the use of a streptavidin–biotin–peroxidase complex (Vector Laboratories, Burlingame, CA). Hematoxylin was used as a counterstain to highlight the nuclei and overall tissue architecture.

### Immunofluorescence

ALCL cells (150,000 per well) were seeded overnight in FCS-free medium on Poly-L-Lysine–coated 8-well chamber slides (Ibidi, Gräfelfing, Germany #80827). Cells were fixed with 4% paraformaldehyde (Thermo Fisher, #043368.9M), permeabilized with 0.5% Triton X-100, then washed with 0.1% Tween-20 in PBS. Non-specific binding was blocked using 10% FCS. Cells were incubated with rabbit anti-ZBTB46 antibody (1:100; Atlas, Stockholm, Sweden, #HPA103997, RRID:AB_1858961) in 2% FCS/PBS, followed by a 2-h incubation at RT with Alexa Fluor 568-conjugated goat anti-rabbit secondary antibody (1:2000; Thermo Fisher, #A11011, RRID:AB_143157) in 2% FCS/PBS. After washing, slides were mounted with DAPI-containing medium (Thermo Fisher, #S36920). Imaging was performed using a Zeiss LSM 880 confocal microscope in Airyscan mode with Zen Black software.

### MicroRNA detection

Retro transcription was performed from 10 µg total RNA using miRCURY LNA RT Kit (Qiagen, Les Ulis, France # 339340), following manufacturer’s instructions. qPCR quantification was performed using miRCURY LNA SYBR Green PCR Kit (Qiagen #339345) following manufacturer’s instructions. SNORD44 RNA was used as housekeeping gene. Probes were listed in **Supplementary Table 1.**

### Apoptosis assay

Apoptosis was detected via a Pacific Blue™ Annexin V kit (BioLegend #640918) or an APC/Fire™ 750 Annexin V (BioLegend, San Diego, CA #640953) following the manufacturer’s instructions. Flow cytometry analysis was performed on a MACSQuant® (Miltenyi Biotec, Paris, France) Analyzer 10 flow cytometer, and the data were analyzed with FlowJo software(BD Biosciences, San Jose, CA).

### Cell cycle analysis

Cell cycle analysis was performed via an EdU Click-iT kit (Invitrogen, Waltham, MA #C10337) following the manufacturer’s instructions.

## Acknowledgments

The study was supported by INSERM, Labex Toucan, Association Eva pour la vie, Féderation Grandir sans Cancer, RACCE Lions Club de Lourdes, the French Ministry of Health and the French National Cancer Institute (CircOma, PRT-K 2022-184), Fondation ARC et Fondation pour la Recherche Médicale supported by its sponsor Nagui and his production company, Banijay Production (EQU202403018019). All grants were obtained by F.M. E.A. and were supported by Labex Toucan/Laboratoire d’Excellence Toulouse Cancer, the association Cassandra, and Labex Toucan/Laboratoire d’Excellence Toulouse Cancer, EUR CARe N°ANR-18-EURE-0003 in the framework of the Programme des Investissements d’Avenir, L.B. and C.B. were supported by the Fondation de France. L.B. was supported by Fondation l’Oréal for Women in Science, Prolific Graine de Chercheur and Association Ellye.

L.C. and S.F. were supported by a grant from the French Ministry of Health and the French National Cancer Institute (CircOma, PRT-K 2022--184). S.F. was supported by a grant from the Deutsche Forschungsgemeinschaft (DFG, grant no. 439441203). S.F. is a participant in the BIH-Charité Clinician Scientist Programme funded by the Charité - Universitätsmedizin Berlin and the Berlin Institute of Health. C.P. and I.D. were supported by FONROGA. The authors thank the staff of the animal facility of UMS US006/CREFRE (Toulouse, France) for their technical assistance. This work benefited from the equipment and services of the iGenSeq core facility (genotyping and sequencing) at the ICM. Part of this work was carried out by the Paris Brain Institute (ICM) Data Analysis Core (DAC) plateform (RRID:SCR_026138, https://dac.institutducerveau.org/). We gratefully acknowledge Thomas Gareau for assistance with RNAseq data analysis.

The study was conducted in accordance with the guidelines of the Declaration of Helsinki and approved by the Institutional Review Board of the Toulouse Hospital Biobank (DC-2020-4074; AC-2020-4031, 2020).

Medical writing for this manuscript was assisted by MPIYP (MC Béné), Paris, France.

## Data availability

The ribo zero full RNA sequencing, the long-read Oxford Nanopore RNA sequencing and STAT3 ChIP-seq datasets are available in the GEO repository (GSE160123, GSE197872 and GSE117164 respectively). The RNA-sequencing datasets generated are available from the corresponding author on reasonable request.

## Author contributions

E.A., L.B. and F.M. designed the research study. E.A., S.F., C.P., I.D., C.Q., R.F., A.Z. and L.B. performed the experiments. E.A., L.B., S.F., C.Q. and F.M. analyzed the data. L.L. and C.Q. were involved in the diagnosis of ALCL. C.B., L.B. and C.G. performed the bioinformatic analysis. S.P. and M.B. helped with discussions and critical reading. L.B. and F.M. wrote the manuscript. All authors have read and agreed to the published version of the manuscript.

