## Supplemental Data for "CircZBTB46, a promising therapeutic target in crizotinib resistant ALK-positive T lymphomas"

**Supplementary Data**

**Supplementary Figure legends**

**Supplementary Figure 1. Expression of circZBTB46, *ZBTB46* and ALK in ALK(+) lymphoma cells and controls.** ZBTB46 and ALK protein expression in ALK(+) ALCL cell lines (KARPAS-299, SU-DHL-1, SUPM2 and COST), MAC2A ALK(-) ALCL cell line and CD3(+) stimulated lymphocytes (CD3 S) was assessed via Western blot analysis. Actin was used as a loading control.

**Supplementary Figure 2. Regulation of *ZBTB46* expression by NPM1-ALK and STAT3 signaling. (A)** Relative expression of *ZBTB46* mRNA, as determined by RNA-Seq, in primary human CD4 T lymphocytes transduced with a lentiviral vector encoding NPM1-ALK [25] and **(B)** *ZBTB46* mRNA expression in human T cells engineered to carry the canonical t(2;5)(p23;q35) translocation by CRISPR/Cas9 genome editing [18]. **(C)** STAT3 chromatin enrichment at the *IRF4*, *STAT3* and *ZBTB46* loci in JB6 and SUDHL1 ALCL cell lines as assessed by *STAT3* ChIP-seq (GSE117164; [26]).

**Supplementary Figure 3. Characterization of *ZBTB46* CRISPR/Cas9 knockout models**. **(A)** PCR-based detection of *ZBTB46* exon 2-5 (Del2-5) and exon 4-5 (Del4-5) deletions at the DNA level. Amplification of wild-type (WT) exons 2, 4 and 5 was used as a control. **(B)** ZBTB46 protein expression in COST wild-type (WT), Del2-5 and Del4-5 clones as assessed by Western blot using an antibody directed against an epitope encoded by exon 2. Actin was used as a loading control. **(C)** Immunofluorescence staining of the ZBTB46 protein in COST WT, Del4-5 and Del2-5 clones. Original magnification, 63x **(D)** Cell cycle distribution determined by EdU incorporation and flow cytometry analysis of COST WT, Del2-5 and Del4-5 clones.

**Supplementary Figure 4. ZBTB46 protein expression after circZBTB46 was targeted via the Cas13 system.** ZBTB46 protein levels in COST, SUPM2 and Pio ALK(+) ALCL cell lines transduced with Cas13 and either control gRNA (CTL) or gRNA targeting circZBTB46, as assessed by Western blot analysis. GAPDH was used as a loading control.

**Supplementary Figure 5. MiR-25-3p regulates PIP5K1C expression in crizotinib-resistant ALK(+) ALCL cells. (A)** PIP5K1C protein levels detected by Western blot in stable COSTR200 cells expressing CRISPRi constructs with control gRNA (CTL) or gRNA targeting the PIP5K1C promoter. **(B)** Cell viability of crizotinib-resistant COSTR200 cells transduced with CRISPRi/gRNA targeting PIP5K1C, followed by crizotinib treatment (1,000 nM, 48 h), as measured by Annexin V/PI staining (flow cytometry). **(C-D)** MicroRNAs were predicted to target the PIP5K1C 3'UTR via **(C)** TargetScanHuman or **(D)** miRTargetLink 2.0. **(E-F)** Crizotinib-resistant ALK(+) ALCL cells (COSTR200) were transfected miR_25-3p mimic (mimic-miR25-3p), a miR-25-3p inhibitor (inhibit-miR25-3p), or the corresponding negative controls (mimic-CTL and inhibit-CTL)**. (E) R**elative expression of miR_25-3p in COSTR200 cells after transfection measured by RT-qPCR. Snord44 was used as an internal control. Data are shown as 2-ΔΔCt relative values. **(F)** Western blot analysis of PIP5K1C protein levels in COSTR200 cells transfected with a miR_25-3p mimic, miR_25-3p inhibitor (inhib), or their respective negative controls. GAPDH served as a loading control. Experiments were performed in triplicate. Data are presented as mean ± SEM. Statistical significance was determined using an unpaired two-tailed Student's t test with Welch's correction**:** $P < 0.05$ (*), $P < 0.001$ (**), ns = not significant.

**Supplementary Figure 6. Tumor growth after PIP5K1C inhibition**

Tumor growth in NSG mice (n=8) injected subcutaneously with COSTR200 cells transduced with PIP5K1C CRISPRi or control CRISPRi and treated with crizotinib (100 mg/kg/day). Tumor volume was monitored over time via calipers and is expressed as the mean ± SEM.

**Supplementary Figure 7. Expression of the *ZBTB46* transcript and circZBTB46 in ALK(+).** Each dot represents normalized RNA-seq expression (Kmer count per billion [$\log_2$(count+1)]) of the indicated RNA in primary biopsy samples, including ALK(+) ALCL ($n$ = 39) and AML samples (IUCT-AML cohort ($n$ = 40; [11,12], reactive lymph nodes as healthy lymphoid tissue) (RLN, $n$ = 9), CD3(+) stimulated (CD3-S, n = 3) or not lymphocytes (CD3-NS, n = 3) and established cell lines (AML, $n$ = 5; ALK(+) ALCL, $n$ = 7, including in house NPM1-ALK–transformed and immortalized T-cell models (M1: lentiviral NPM1-ALK(+) [13]; 673: CRISPR-Cas9 NPM1-ALK(+ )[1]; see Supplementary Figure 2). Statistical significance was assessed via an unpaired two-tailed Student's t test with Welch's correction: $P < 0.05$ (*), $P < 0.01$ (**), $P < 0.001$ (***), ns = not significant. Data are expressed as the means ± SEMs.

*Babin et al, Supplementary Figure 1*

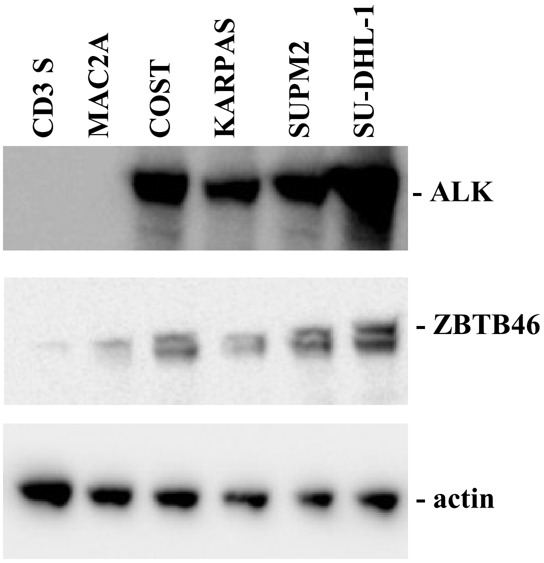

*Babin et al, Supplementary Figure 2*

A

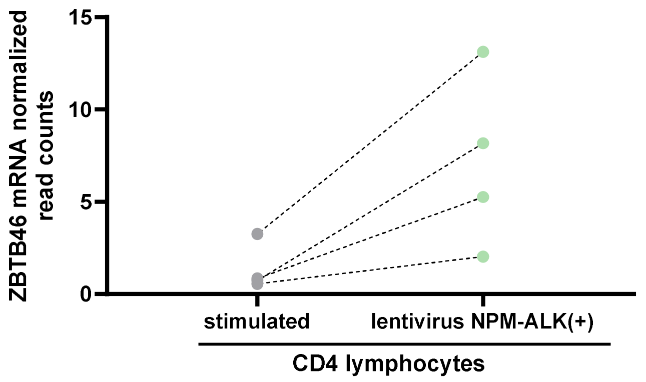


B

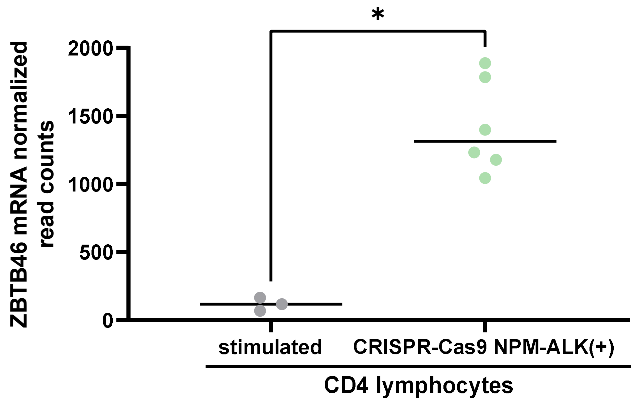


C

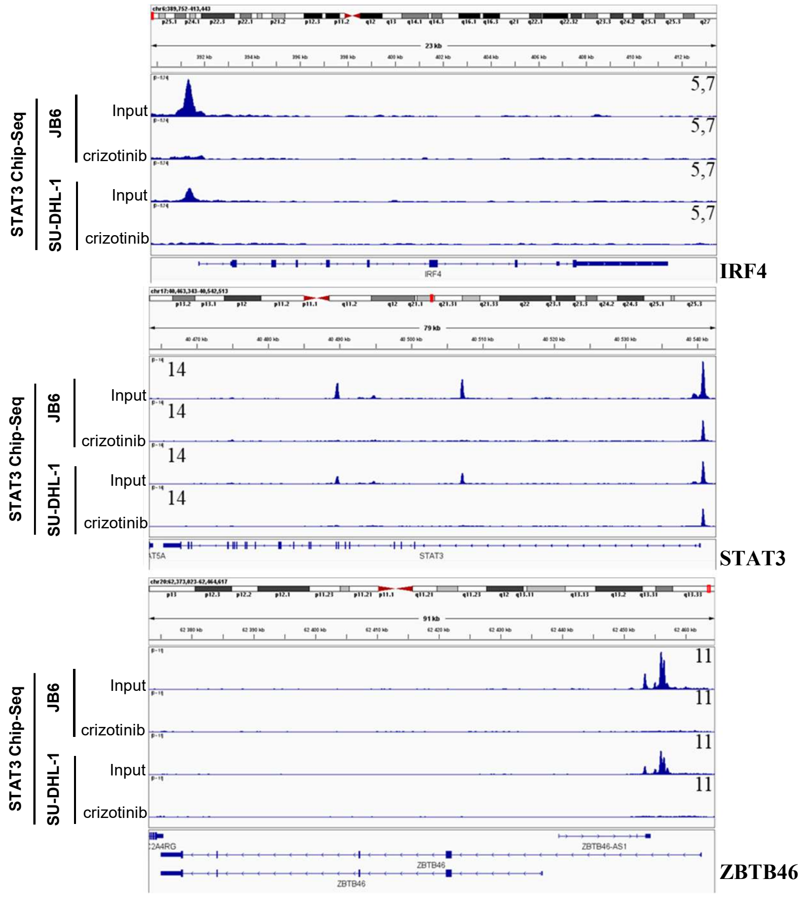

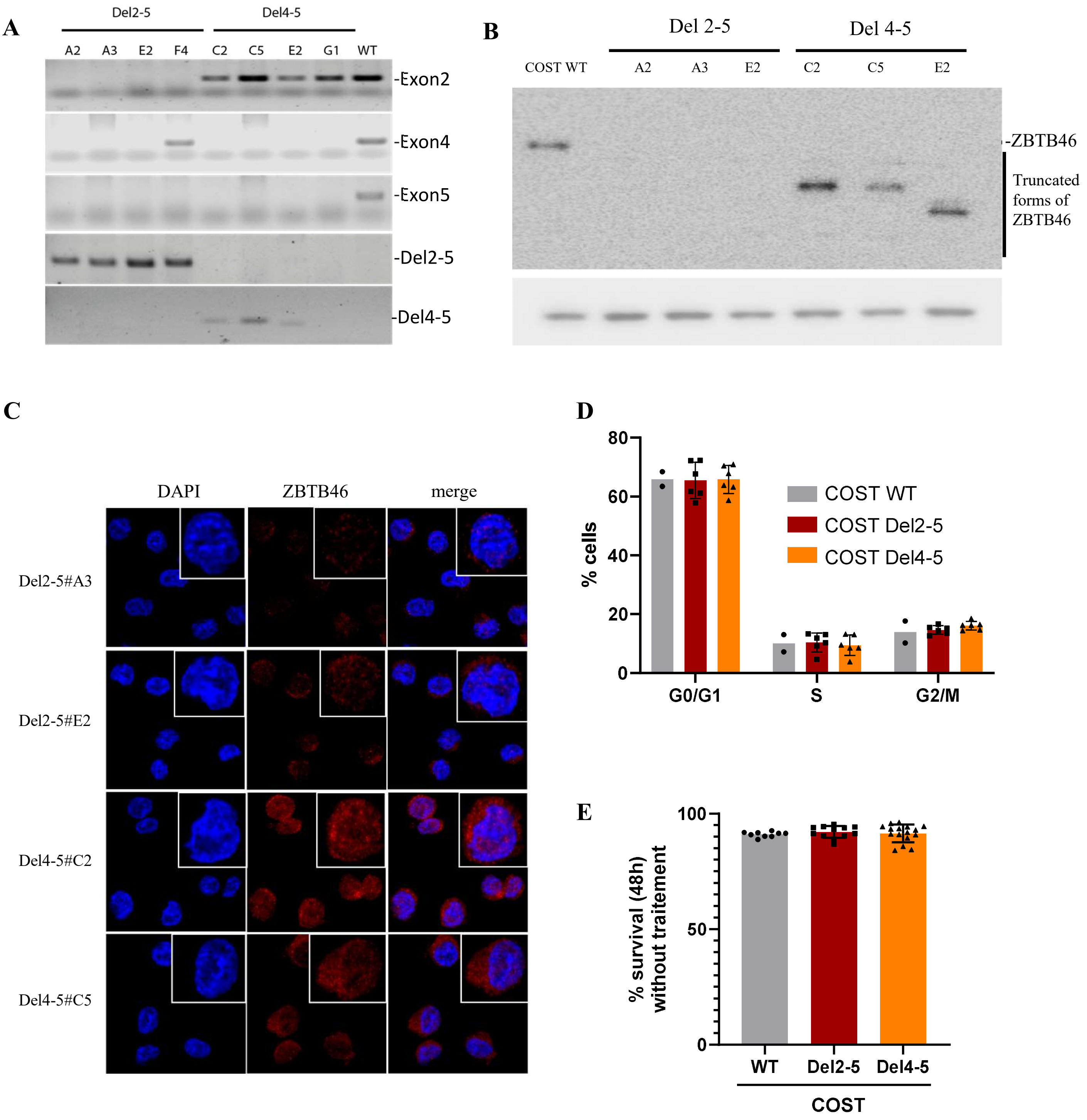
A
Del2-5
Del4-5
A2
A3
E2
F4
C2
C5
E2
G1
WT
-Exon2
-Exon4
-Exon5
-Del2-5
-Del4-5
B
Del 2-5
Del 4-5
COST WT
A2
A3
E2
C2
C5
E2
-ZBTB46
Truncated forms of ZBTB46
C
DAPI
ZBTB46
merge
Del2-5#A3
Del2-5#E2
Del4-5#C2
Del4-5#C5
D
% cells
80
60
40
20
0
G0/G1
S
G2/M
COST WT
COST Del2-5
COST Del4-5
E
% survival (48h) without traitement
100
50
0
WT
Del2-5
Del4-5
COST

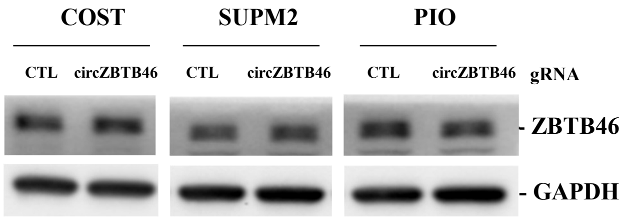
COST
SUPM2
PIO
CTL
circZBTB46
CTL
circZBTB46
CTL
circZBTB46
gRNA
- ZBTB46
- GAPDH

A

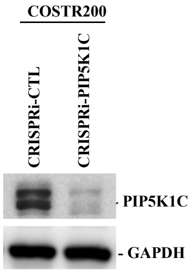
COSTR200
CRISPRi-CTL
CRISPRi-PIP5K1C
PIP5K1C
GAPDH

B

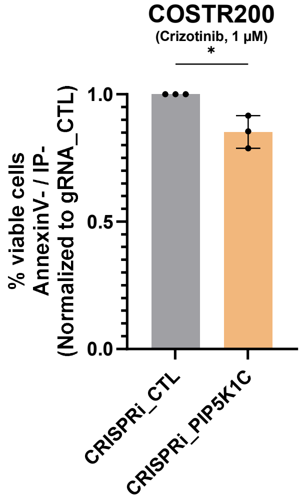
COSTR200
(Crizotinib, 1 µM)
*
% viable cells
AnnexinV- / IP-
(Normalized to gRNA_CTL)
1.0
0.5
0.0
CRISPRi_CTL
CRISPRi_PIP5K1C

C

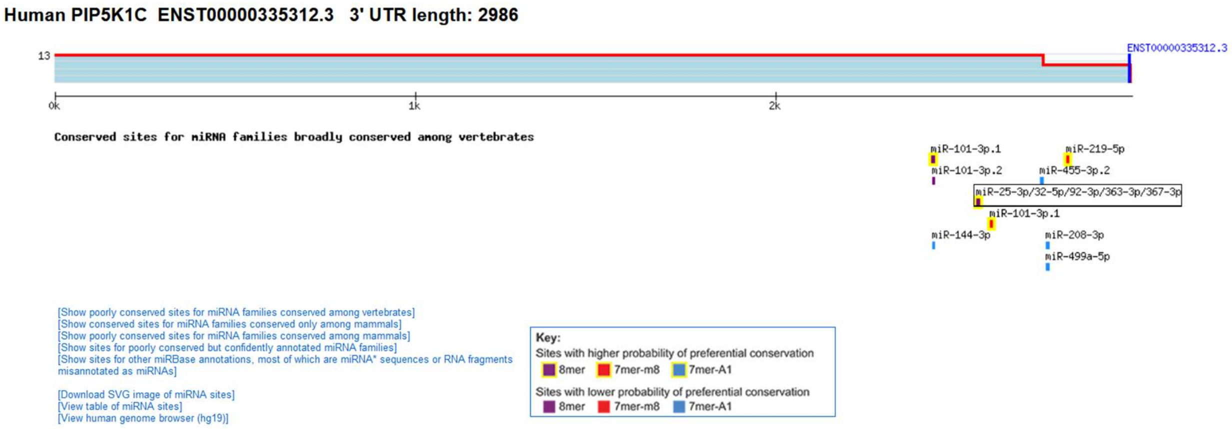
Human PIP5K1C ENST00000335312.3 3' UTR length: 2986
13
ENST00000335312.3
0k
1k
2k
Conserved sites for miRNA families broadly conserved among vertebrates
miR-101-3p.1
miR-219-5p
miR-101-3p.2
miR-455-3p.2
miR-25-3p/32-5p/92-3p/363-3p/367-3p
miR-101-3p.1
miR-144-3p
miR-208-3p
miR-499a-5p
[Show poorly conserved sites for miRNA families conserved among vertebrates]
[Show conserved sites for miRNA families conserved only among mammals]
[Show poorly conserved sites for miRNA families conserved among mammals]
[Show sites for poorly conserved but confidently annotated miRNA families]
[Show sites for other miRBase annotations, most of which are miRNA* sequences or RNA fragments misannotated as miRNAs]
[Download SVG image of miRNA sites]
[View table of miRNA sites]
[View human genome browser (hg19)]
Key:
Sites with higher probability of preferential conservation
8mer
7mer-m8
7mer-A1
Sites with lower probability of preferential conservation
8mer
7mer-m8
7mer-A1

D

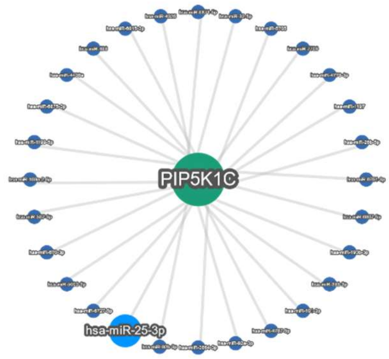
PIP5K1C
hsa-miR-25-3p

E

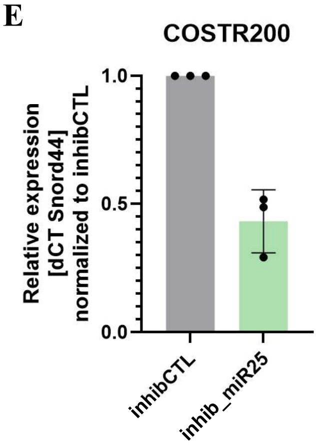
COSTR200
Relative expression
[dCT Snord44]
normalized to inhibCTL
1.0
0.5
0.0
inhibCTL
inhib_miR25

F

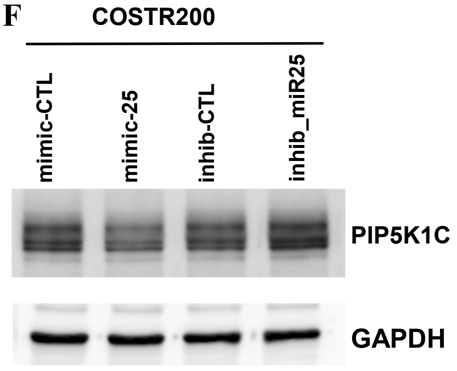
COSTR200
mimic-CTL
mimic-25
inhib-CTL
inhib_miR25
PIP5K1C
GAPDH

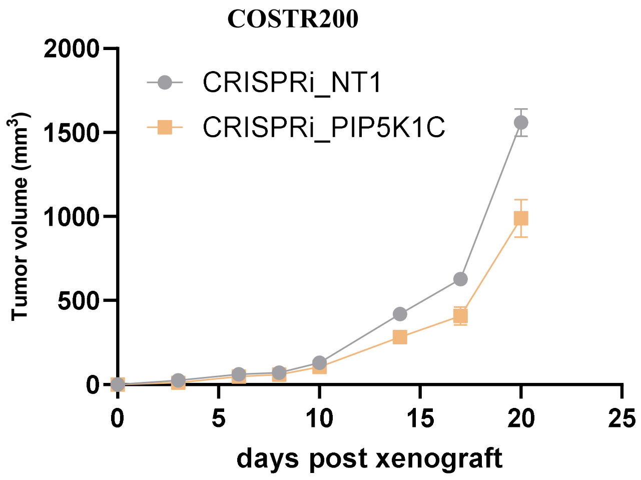
COSTR200
CRISPRi_NT1
CRISPRi_PIP5K1C
Tumor volume (mm3)
2000
1500
1000
500
0
0
5
10
15
20
25
days post xenograft

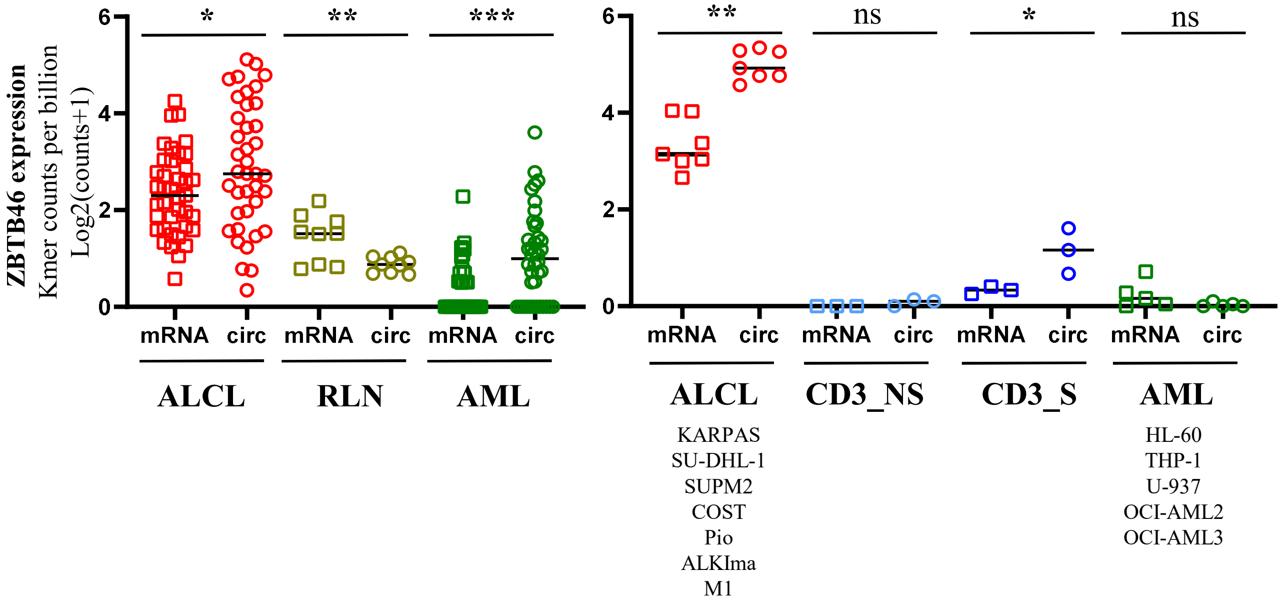
ZBTB46 expression
Kmer counts per billion
Log2(counts+1)
6
4
2
0
*
**
***
mRNA circ mRNA circ mRNA circ
ALCL
RLN
AML
6
4
2
0
**
ns
*
ns
mRNA circ mRNA circ mRNA circ mRNA circ
ALCL
CD3_NS
CD3_S
AML
KARPAS
SU-DHL-1
SUPM2
COST
Pio
ALKIma
M1
HL-60
THP-1
U-937
OCI-AML2
OCI-AML3

**Supplementary Table 1**

| | |
|---|---|
| **Primers sequences** | |
| **RNA** | |
| PIP5K1C-F | GGCAGGTTTGGCTCAGAAGA |
| PIP5K1C-B | CCTCGATGGCCCAACTTCTT |
| STAT3-F | TGCCTGTGGGAAGAATCACG |
| STAT3-B | TCTGCTGCTTCTCCGTCACC |
| MLN51-F | TAATCCCAGTTACCCTTATGCTCCA |
| MLN51-B | GTTATAGTAGGTCACTCCTCCATATACCTGT |
| ABL-F | TGGAGATAACACTCTAAGCATAACTAAAGGT |
| ABL-B | GATGTAGTTGCTTGGGACCCA |
| ZBTB46-F | TGCGAGATCTGCGGGAAGAA |
| ZBTB46-B | GCACACCTTGCACACATACTT |
| circZBTB46-F | CCGGTAGTGGGACGTGATTT |
| circZBTB46-B | ACTCGCTGTCCCAGTCTGTA |

| | |
|---|---|
| **DNA** | |
| ChIP_ZBTB46-F1 | CACAAAAGCCCTGAACCACG |
| ChIP_ZBTB46-B1 | AGGCGGTAAGTGTTCCTTGG |
| ChIP_ZBTB46-F2 | ATCAAACCCTTCCTGGCTGG |
| ChIP_ZBTB46-B2 | TTCATCATGTTGGCCAGGCT |
| ChIP_promSTAT3-F | TGATTCCCGCGTGGTAAGAG |
| ChIP_promSTAT3-B | ACCGGAATGTCCTGCTGAAA |
| ChIP_promGAPDH-F | GCACGGAAGGTCACGATGT |
| ChIP_promGAPDH-B | CCGGGATTGTCTGCCCTAAT |
| ChIP_promALK-F | GCAGAACAACTCTGCCAGAC |
| ChIP_promALK-B | GCATTCCTGGGAGCTTAGGA |
| ZBTB46-Exon4-F | GTACATCCCAAGCCCTGCT |
| ZBTB46-Exon4-B | GTGAACTTCTTCCCGCAGAT |
| ZBTB46-Exon2-F | GTAGAAGAGGCGACACCAGG |
| ZBTB46-Exon2-B | TTGAAGACCTTGCCCTCCAC |
| ZBTB46-Exon5-F | CCCTATCTGGAGGACCCTGA |
| ZBTB46-Exon5-B | TAGGAGAGCCAGGCGAAGT |

| | |
|---|---|
| **siRNA sequences** | |
| circZBTB46_siRNA_1 | CACUCGCUGUCCCAGUCUGUA |
| circZBTB46_siRNA_2 | UCGCUGUCCCAGUCUGUAGAA |
| CTL_siRNA | UAAUGUAUUGGAACGCAUA |
| NPM-ALK_siRNA | GGGCGAGCUACUAUAGAAA |
| SmartPool-ON-TARGETplus Human STAT3 siRNA Dharmacon | L-003544-00-0020 |

| | |
|---|---|
| **shRNA cloning sequences** | |
| sh_circZBTB46_1_S | CCGGTACAGACTGGGACAGCGAGTGCTCGAGCACTCGCTGTCCCAGTCTGTATTTTTG |
| sh_circZBTB46_1_AS | AATTCAAAAATACAGACTGGGACAGCGAGTGCTCGAGCACTCGCTGTCCCAGTCTGTA |

| | |
|---|---|
| **guide RNA sequences** | |
| **Cas13** | |
| Cas13gRNA-circZBTB46 | TTCTACAGACTGGGACAGCGAGT |
| Cas13gRNA-CTL | TAGATTGCTGTTCTACCAAGTAATCCAT |
| **Cas9** | |
| gRNA-ZBTB46-Exon2 | TGGAAATCACGTCCCACTAC |
| gRNA-ZBTB46-Exon4 | TGAATGAGTTCACGGTGATC |
| gRNA-ZBTB46-Exon5 | GTTGGCGGACGACAAGGATG |
| **CRISPRi** | |
| gRNA_PIP5K1C_prom | ATCTTACTGACCTAATACGA |
| gRNA_CTL | CTGAAAAAGGAAGGAGTTGA |

| | |
|---|---|
| **idRIP probes sequences** | |
| 3_bioteg_circZBTB46 | TTCTACAGACTGGGACAGCGA |
| 3_bioteg_CTL | TAATGTATTGGAACGCATA |

| | |
|---|---|
| **mimic sequences** | |
| hsa-miR-25-3p | 5' phosphate - UCAGACCGAGACAAGUGCAAUG dTdT |
| control | 5' phosphate - UAAUGUAUUGGAACGCAUA dTdT |

| | |
|---|---|
| **Probes for microRNA detection by qPCR** | |
| hsa-miR-25-3p miRCURY LNA miRNA PCR Assay | GeneGlobe ID: YP00204361 |
| SNORD44 (hsa) miRCURY LNA miRNA PCR Assay | GeneGlobe ID: YP00203902 |

### Supplementary Table 2

AGTCTGTAGAAGAGGCGACACCAGGGCTTCCAAATGAACAACCGAAAGGAAGATATGGAAATCACGTCCCACTACCGGCA
CCTGCTGCGGGAGCTCAACGAGCAGAGGCAGCACGGCGTCCTGTGCGACGTCTGCGTGGTCGTGGAGGGCAAGGTCTTCA
AGGCGCACAAGAACGTCCTGCTGGGCAGCAGCCGCTACTTCAAGACGCTCTACTGCCAGGTGCAGAAGACGTCGGAGCAG
GCCACGGTCACGCACCTGGACATCGTCACGGCCCAGGGCTTCAAGGCCATCATCGACTTCATGTACTCAGCGCACCTGGC
GCTCACCAGCAGGAACGTCATCGAGGTGATGTCAGCCGCCAGCTTCCTGCAGATGACGGACATCGTGCAAGCCTGCCACG
ACTTCATCAAGGCGGCGCTGGACATCAGCATCAAGTCGGACGCCTCAGATGAGCTTGCGGAGTTCGAGATCGGCGCCTCG
TCCAGCAGCAGCACGGAAGCTCTCATCTCGGCCGTGATGGCTGGGAGGAGCATCTCCCCGTGGCTGGCACGGCGAACGAG
TCCTGCCAATTCTTCCGGAGACTCGGCCATCGCCAGCTGTCACGACGGAGGGAGCAGCTACGGGAAAGAGGATCAGGAGC
CCAAGGCCGATGGCCCTGATGATGTTTCTTCACAGCCTCTATGGCCTGGAGACGTGGGCTACGGGCCTCTGCGCATCAAG
GAAGAGCAGGTTTCACCGTCTCAGTACGGAGGGAGCGAGCTGCCTTCTGCCAAGGACGGTGCAGTACAGAACTCTTTCTC
AGAGCAGAGTGCTGGTGATGCCTGGCAGCCCACGGGCCGAAGGAAGAATCGGAAAAACAAAGAGACCGTCCGGCACATCA
CACAGCAGGTGGAAGATGACAGCCGGGCCAGCTCCCCGGTGCCGTCCTTCCTGCCGACGTCGGGGTGGCCGTTCAGCAGC
CGAGACTCAAATGCGGACCTGTCCGTCACCGAAGCCAGCAGCTCCGACAGCCGAGGAGAGAGGGCCGAGCTCTATGCACA
GGTGGAGGAGGGTCTCCTGGGAGGAGAAGCCAGCTATCTGGGCCCTCCCCTCACCCCAGAGAAGGACGACGCCCTGCATC
AGGCCACCGCGGTGGCCAACCTGCGCGCGGCGCTCATGAGTAAGAACAGCCTGCTGTCGCTGAAGGCCGACGTGCTGGGG
GATGACGGCTCCCTGCTGTTCGAGTACCTGCCCAGAGGGGCCCACTCGCTGTCCC
